# Intraspecific diversity of fission yeast mitochondrial genomes

**DOI:** 10.1101/624742

**Authors:** Yu-Tian Tao, Fang Suo, Sergio Tusso, Yan-Kai Wang, Song Huang, Jochen B. W. Wolf, Li-Lin Du

## Abstract

The fission yeast *Schizosaccharomyces pombe* is an important model organism, but its natural diversity and evolutionary history remain under-studied. In particular, the population genomics of the *S. pombe* mitochondrial genome (mitogenome) has not been thoroughly investigated. Here, we assembled the complete circular-mapping mitogenomes of 192 *S. pombe* isolates *de novo*, and found that these mitogenomes belong to 69 non-identical sequence types ranging from 17618 bp to 26910 bp in length. Using the assembled mitogenomes, we identified 20 errors in the reference mitogenome and discovered two previously unknown mitochondrial introns. Analysing sequence diversity of these 69 types of mitogenomes revealed two highly distinct clades, with only three mitogenomes exhibiting signs of inter-clade recombination. This diversity pattern suggests that currently available *S. pombe* isolates descend from two long-separated ancestral lineages. This conclusion is corroborated by the diversity pattern of the recombination-repressed *K*-region located between donor mating-type loci *mat2* and *mat3* in the nuclear genome. We estimated that the two ancestral *S. pombe* lineages diverged about 31 million generations ago. These findings shed new light on the evolution of *S. pombe* and the datasets generated in this study will facilitate future research on genome evolution.

## INTRODUCTION

The fission yeast *Schizosaccharomyces pombe* is a unicellular fungal species belonging to the Taphrinomycotina subphylum of the Ascomycota phylum (Liu et al. 2009). The first descriptions of this species in the 1890s reported it as a microorganism associated with fermented alcoholic drinks, including its presence in East African millet beer and in the fermenting sugar-cane molasses for making a distilled liquor (Batavia Arrack) in Indonesia (Lindner 1893; Vorderman 1893; Eijkman 1894; Barnett & Lichtenthaler 2001; Lodder & Kreger-Van Rij 1952). Since then, *S. pombe* has been found in various human-associated environments throughout the world, but has never been isolated in truly wild settings (Brown et al. 2011; Jeffares 2018; Jeffares et al. 2015). In 1947, Urs Leupold, the founder of fission yeast genetics, selected an *S. pombe* isolate from French grape juice as the subject of his PhD research, and this strain (hereafter referred to as the Leupold strain) has essentially been the only strain used for modern *S. pombe* molecular biology studies (Osterwalder 1924; Leupold 1950; Hu et al. 2015). In 2002, the complete genome sequence of the Leupold strain was published, making *S. pombe* the sixth eukaryotic species with a sequenced genome (Wood et al. 2002). Today, *S. pombe* is recognized as one of the most prominent model organisms for understanding the molecular mechanisms of cellular processes (Hayles & Nurse 2018; Hoffman et al. 2015).

In recent years, the intraspecific genomic diversity of *S. pombe* has begun to be investigated (Brown et al. 2011; Rhind et al. 2011; Avelar et al. 2013; Clément-Ziza et al. 2014; Fawcett et al. 2014; Zanders et al. 2014; Hu et al. 2015; Jeffares et al. 2015, 2017). In particular, Jeffares et al. sequenced the genomes of 161 *S. pombe* isolates and comprehensively explored genomic variation within this species (Jeffares et al. 2015, 2017). However, the breadth and depth of knowledge on the natural diversity and evolutionary history of *S. pombe* remain limited, especially compared to the other model yeast species, *Saccharomyces cerevisiae* (Duan et al. 2018; Peter et al. 2018).

The mitochondrion originates from a bacterial endosymbiont and, after extensive reductive evolution, still retains a small genome (Lang et al. 1997). Compared to the nuclear genome, the smaller size, higher copy number, and lower level of recombination of the mitochondrial genome (mitogenome) have long made it an attractive subject for intraspecific comparative studies, shedding light on the evolution of many species including humans (Ingman et al. 2000). Recently, the population mitogenomic approach has begun to be applied to fungal species, leading to new insights about population structure and evolutionary dynamics (Freel et al. 2015; Jung et al. 2012; Leducq et al. 2017; Wolters et al. 2015).

The mitogenome of the Leupold strain of *S. pombe* was completely sequenced more than 10 years before its nuclear genome (Lang 1984; Lang et al. 1985, 1987; Trinkl et al. 1989). It contains 2 rRNA genes (*rnl* and *rns*), 8 protein-coding genes (*atp6*, *atp8*, *atp9*, *cob*, *cox1*, *cox2*, *cox3*, and *rps3* which is also known as *var1*), a gene encoding the RNA component of mitochondrial RNaseP (*rnpB*), and 25 tRNA genes (Bullerwell et al. 2003; Schäfer 2003). No complete mitogenome sequences of any other *S. pombe* isolates have been reported thus far. Restriction fragment analysis and limited Sanger sequencing have indicated that presence-absence polymorphisms of mitochondrial introns are widespread among *S. pombe* isolates (Zimmer et al. 1984, 1987), but an accurate and thorough understanding of intraspecific mitogenomic variation of *S. pombe* is still lacking.

In this study, we used both published and newly generated genome sequencing data to perform *de novo* assembly of the mitogenomes in 199 *S. pombe* isolates. We successfully assembled the complete mitogenome sequences of 192 isolates. Analysing these mitogenome sequences led to the discovery of reference mitogenome errors, new mitochondrial introns, and divergence patterns providing new insights into the evolutionary history of *S. pombe*. In particular, we show that *S. pombe* isolates descend from two long-separated ancient lineages. An independent study published after the initial submission of this article reaches the same conclusion through analysing the nuclear genome diversity of *S. pombe* isolates (Tusso et al. 2019).

## MATERIALS AND METHODS

### Previously published genome sequencing data of 161 JB strains

The Illumina genome sequencing data of 161 *S. pombe* strains with names that begin with the initials JB were downloaded from European Nucleotide Archive (ENA) according to the ENA accession numbers given in Supplementary Table 7 of Jeffares et al. (Jeffares et al. 2015), and are listed in Supplementary Table S1. For 11 sequencing runs that belong to the ENA Study accession number PRJEB6284, we noticed discrepancies between the read numbers reported in Supplementary Table 7 of Jeffares et al. 2015 and the read numbers in the data downloaded from ENA. The authors of the prior study confirmed that strain name mix-ups had occurred during the submission of the sequencing data to the ENA by The Genome Analysis Centre (TGAC) (Daniel Jeffares, personal communication). A list of these 11 sequencing runs with their correct corresponding strain names is provided as Supplementary Table S2.

**Table 1.** Differences between the reference *S. pombe* mitogenome (NC_001326.1) and MT1

| Reference genome position | Reference genome sequence | MT1 sequence | Type of sequence difference | Affected gene | Amino acid difference* | Detected before by Iben et al. (Iben et al. 2011) |
| --- | --- | --- | --- | --- | --- | --- |
| 1788 | C | T | single-nucleotide substitution | *rnl* | N/A | Yes |
| 2091 | A | G | single-nucleotide substitution | *rnl* | N/A | Yes |
| 2364 | C | CC | insertion | *rnl* | N/A | No |
| 4102 | C | G | single-nucleotide substitution | *rns* | N/A | Yes |
| 4463 | C | A | single-nucleotide substitution | *rns* | N/A | Yes |
| 4525 | AC | A | deletion | *rns* | N/A | No |
| 4777 | AA | A | deletion | intergenic | N/A | No |
| 6906 | A | T | single-nucleotide substitution | *cox1-l2b* | S21C | Yes |
| 8376 | C | CTAC | insertion | *cox1* | an extra Y residue after Y400 | No |
| 8529 | G | T | single-nucleotide substitution | *cox1* | E451D | Yes |
| 10309 | TC | CT | double-nucleotide substitution | *cob* | I46T | Yes |
| 11218 | T | A | single-nucleotide substitution | *cob-l1* | I121N | Yes |
| 11644 | A | C | single-nucleotide substitution | *cob-l1* | H263P | Yes |
| 13196 | C | T | single-nucleotide substitution | *cob-l1* | synonymous | Yes |
| 13751 | C | T | single-nucleotide substitution | *cob* | synonymous | Yes |
| 15081 | TA | T | deletion | *atp6* | L109F and R111G | Yes |
| 15088 | G | GA | insertion | *atp6* |  | Yes |
| 15200 | G | T | single-nucleotide substitution | *atp6* | V149F | Yes |
| 18648 | G | A | single-nucleotide substitution | *cox2* | V30I | Yes |
| 19152 | A | G | single-nucleotide substitution | *cox2* | S198G | Yes |

\*Abbreviation: N/A, not applicable.

### Genome sequencing of 38 strains from culture collections in China and USA

To explore intraspecific diversity beyond the previously analysed strains, we acquired 38 *S. pombe* strains from four culture collections in China and one culture collection in the USA: 20 strains from CGMCC (China General Microbiological Culture Collection Center), 13 strains from CICC (China Center of Industrial Culture Collection), 1 strain from CICIM (Culture and Information Centre of Industrial Microorganisms of China Universities), 1 strain from CFCC (China Forest Culture Collection Center), and 3 strains from NRRL (United States Department of Agriculture Agricultural Research Service Culture Collection) (Supplementary Table S1). Only three of these 38 strains have isolation information: CGMCC 2.1043 was isolated from fermented grains for making Moutai, a Chinese liquor; NRRL Y-11791 was from reconstituted lime juice (location unknown); NRRL Y-48646 was from a wine producing company (location unknown). Single-cell-derived clones of these strains were deposited into our laboratory strain collection (DY collection) and given strain names that begin with the initials DY (Supplementary Table S1). Cells grown on YES solid media were used for genomic DNA preparation using the MasterPure Yeast DNA Purification Kit (Epicentre). The kit manufacturer’s protocol was followed, with the exception of lysing the cells by glass bead beating in a FastPrep-24 homogenizer (MP Biomedicals) for 20 seconds at a speed setting of 6.4 m/s. The sequencing library for DY15505 was constructed using the NEBNext DNA Library Prep Master Mix (NEB). For the other 37 strains, tagmentation-based sequencing library preparation was performed using home-made Tn5 transposase (Picelli et al. 2014). Post-tagmentation gap filling and PCR amplification were performed using the KAPA HiFi HotStart PCR Kit (Kapa Biosystems) with the following cycling parameters: 3 min at 72°C, 30 sec at 95°C, and then 11 cycles of 10 sec at 95°C, 30 sec at 55°C, and 30 sec at 72°C. AMPure XP beads (Beckman Coulter) were used to select PCR product in the size range of 400 bp to 700 bp. Paired-end sequencing was performed using Illumina HiSeq 2000 (2×96 read pairs), HiSeq 2500 (2×100 read pairs or 2×101 read pairs), or HiSeq X Ten sequencers (2×150 read pairs). Sequencing data for these 38 strains have been deposited at NCBI SRA under accession numbers SRR8698890–SRR8698927.

### *De novo* assembly of the mitogenomes

The genome sequencing data for the above mentioned 199 strains (161 JB strains and 38 DY strains) were cleaned by Trimmomatic version 0.32 with options LEADING:30, TRAILING:30, SLIDINGWINDOW:4:30, and MINLEN:80 (MINLEN:130 for 150-bp HiSeq X Ten reads) (Bolger et al. 2014). We found empirically that *de novo* assembly of the mitogenome requires data downsampling with 300,000 cleaned read pairs being a suitable downsampling target data size except for longer-read-length HiSeq X Ten-generated data, which required a lower downsampling target read number. Downsampling was performed using the software seqtk (https://github.com/lh3/seqtk). Three independent sets of randomly downsampled data were obtained using the seed numbers 100, 500, and 800. *De novo* assembly was performed using A5-miseq version 20150522 (Coil et al. 2015). From the A5-miseq output we selected the mtDNA-containing contigs based on length and sequence. Custom Perl scripts were used to trim overlapping sequences at the end of the mtDNA contigs and set the starting position of the circular-mapping mtDNA to that of the reference *S. pombe* mitogenome (accession number NC_001326.1). Full-length mitogenome assemblies were usually obtained from all three sets of downsampled data. Pilon version 1.21 was used to polish the assemblies (Walker et al. 2014). Polished assemblies were verified by mapping all cleaned reads of a strain to the corresponding assembly and manually examining the mapping results on a genome browser. No assembly errors were apparent during manual inspection, nor was any obvious heteroplasmy observed. In total, we obtained full-length mitogenome assemblies for 192 of the 199 strains (Supplementary Table S1). Among these, there were 69 different types of mitogenome sequences which we designated as MT type 1 to 69 or, for brevity, MT1 to MT69 (Supplementary Table S1).

### Validation of the mitogenome assemblies using third-generation sequencing

To validate the assemblies based on Illumina-generated short reads, we assembled mitogenomes using long read data from two types of third-generation sequencing technology: the RSII platform of Pacific Biosciences (PacBio) and MinIOn by Oxford Nanopore (Tusso et al. 2019) (data deposited in Bioproject PRJNA527756 of the sequencing read archive at the National Center for Biotechnology Information). PacBio data for 15 strains (JB4, JB22, JB760, JB842, JB853, JB858, JB872, JB873, JB900, JB918, JB934, JB939, JB1197, JB1205, and JB1206) and MinION data for 7 strains (JB22, JB760, JB858, JB873, JB934, JB1197, and JB1205) were used. *De novo* assembly was performed using Canu 1.5 (Koren et al. 2017) with default parameters followed by polishing using the package BridgeMapper of the SMRT Analysis Software v2.3.0. A second polishing was performed using short reads aligned with BWA 0.7.15 and Pilon 1.22 (Walker et al. 2014). Customised python scripts were used to identify and trim the resulting mitogenome assemblies (available at https://github.com/EvoBioWolf/SchPom_mito/). For comparison, long-read-based mitogenome assemblies were aligned to the corresponding short-read-based ones using MAFFT 7.407 (Katoh & Standley 2013). Because the mitogenome of JB842 was not assembled from short reads, PacBio-based assembly of JB842 was compared to MT47, the short-read-based assembly of JB851 and JB857, which share the same nuclear genome type with JB842. In all cases, the long-read-based assemblies were identical to the corresponding short-read-based assemblies.

### Mitogenome annotation

Protein-coding genes in the assembled mitogenomes were annotated using MFannot based on genetic code 4 (the only difference between genetic code 4 and the standard code is UGA being a tryptophan codon, not a stop codon) (Lang et al. 2007; Valach et al. 2014). MFannot was also used for predicting intronic regions and intron types (group I or group II intron). tRNA and rRNA annotations were transferred from the reference mitogenome using the software RATT (Otto et al. 2011). The EMBL-format reference annotation file required by RATT was generated from the GenBank format file using the software Artemis (Carver et al. 2012), which was also used to convert the RATT output from EMBL format to GenBank format. Results of software-based annotation were verified by manual inspection. The annotations of mt-tRNAArg(UCU) and mt-tRNAGlu(UUC) were revised according to the recently published *S. pombe* mitochondrial transcriptome analysis (the starting and ending positions of the former were shifted upstream for one nucleotide and six nucleotides, respectively, and the ending position of the latter was shifted downstream for one nucleotide) (Shang et al. 2018). The 69 types of mitogenomes (MT1–MT69) together with their annotations have been deposited at GenBank under accession numbers MK618072– MK618140. The lengths of these mitogenomes, the total lengths of different types of sequence features in these mitogenomes, and the intron presence-absence patterns are listed in Supplementary Table S3.

Published sequences and annotations of the mitogenomes of the three other *Schizosaccharomyces* species were used for the analysis of genes in these three mitogenomes (accession numbers NC_004312.1 and AF275271.2 for *S. octosporus*, accession numbers NC_004332.1 and AF547983.1 for *S. japonicus*, and accession number MK457734 for *S. cryophilus*) (Bullerwell et al. 2003; Rhind et al. 2011).

### *De novo* assembly of the recombination-repressed *K*-region in the nuclear genome

Using the same set of Illumina genome sequencing data of 199 *S. pombe* strains, we performed targeted *de novo* assembly of the *K*-region. For this purpose, we employed the assembler software TASR version 1.6.2 in the *de novo* assembly mode (-i 1 mode) (https://github.com/warrenlr/TASR) (Warren & Holt 2011). Because the 4.3-kb centromere-repeat-like *cenH* element within the *K*-region cannot be assembled using short sequencing reads, we chose the 4.6-kb reference genome sequence of the *mat2*–*cenH* interval (nucleotides 4557–9169 of GenBank accession number FP565355.1) and the 1.9-kb reference genome sequence of the *cenH*–*mat3* interval (nucleotides 13496–15416 of GenBank accession number FP565355.1) as the input sequences provided to TASR for read recruitment. Based on read mapping, 12 of the 199 strains lacked the *K*-region (Supplementary Table S1). These 12 strains included JB22 (Leupold’s 972 strain), an *h^−S^* mating type strain in which the *K*-region is known to be absent (Beach & Klar 1984). For 150 (80%) of the remaining 187 strains, we were able to generate complete assemblies corresponding to the two target sequences (Supplementary Table S1). The failure to fully assemble the sequences for the other 37 strains appeared to be mainly owing to insufficient sequencing depth, as 92% (110/119) of the strains with >40× average nuclear genome sequencing depth (based on cleaned reads) had fully assembled sequences, whereas only 59% (40/68) of the strains with <40× sequencing depth had fully assembled sequences (Supplementary Table S1). We concatenated the fully assembled *mat2*–*cenH* interval, 100 Ns (representing the unassembled *cenH* sequence), and the fully assembled *cenH–mat3* interval together as the *K*-region sequence. Mapping reads to the assembled *K*-region sequences showed that, for strains that appear to have more than one copy of the *K*-region based on read depth (Supplementary Table S1), no obvious sequence differences exist between *K*-region copies, except for one single-nucleotide inter-copy variation in DY29155 and DY29156. Among the *K*-region sequences of 150 strains, there are 29 non-identical sequence types. We designated them *K*-region type 1 to 29 or, for brevity, K1 to K29 (Supplementary Table S1). In particular, we assigned *K*-region type 1 (K1) to the *K*-region sequence in JB50 (Leupold’s 968 *h^90^* strain), a strain that should have the same *K*-region sequence as the reference genome. However, K1 differed in 34 positions from the *K*-region sequence in the reference genome, including 26 nucleotide substitutions and 8 one-base indels. For all but one of these 34 positions, the reference genome alleles did not exist in any *K*-region types, whereas the alleles of K1 were shared by other *K*-region types. Thus, these differences are most likely due to reference sequence errors. The 29 types of *K*-region sequences (K1–K29) have been deposited at GenBank under accession numbers MK618141–MK618169.

### Phylogenetic tree construction

We used two methods to construct phylogenetic trees based on gene sequences present in all 69 MT types. In the first method, we used the non-intronic nucleotide sequences of 9 genes (*rnl*, *rns*, *cox1*, *cox3*, *cob*, *atp6*, *atp8*, *atp9*, and *cox2*) present in the mitogenomes of all four fission yeast species to construct a neighbour-joining tree based on the p-distance model in MEGA 7.0.18 (Kumar et al. 2016). Bootstrap analysis with 1000 replicates was performed. In the second method, we employed MEGA to construct a maximum likelihood tree using the non-intronic nucleotide sequences of the above 9 genes plus *rps3* and *rnpB*. The model recommended by MEGA, TN93+G+I, was used. Bootstrap analysis with 1000 bootstrap replicates was performed. For the construction of the phylogenetic trees of introns, a maximum likelihood tree of each intron was constructed with 100 bootstrap replicates using the model suggested by MEGA. For the construction of the phylogenetic trees of intron-encoded proteins (IEPs), we obtained protein sequences closely related to *Schizosaccharomyces* IEPs by BLASTP search of NCBI nr database. Maximum likelihood trees were constructed with 100 bootstrap replicates using the model suggested by MEGA. A neighbour-joining tree and a maximum likelihood tree of the *K*-region were also constructed using MEGA.

### ADMIXTURE analysis and heatmap analysis

For the 69 MT types, nucleotide substitution variants were identified from the sequence alignment of intron-removed sequences. Bi-allelic single-nucleotide variants (SNVs) were merged into bi-allelic multi-nucleotide variants (MNVs) if two neighbouring bi-allelic SNVs were less than 15 bp apart and shared the same allelic partition of the 69 MT types. We used custom Perl scripts and PLINK 1.07 to generate a binary PLINK BED format file, which was used as input for ADMIXTURE version 1.3.0 (Alexander et al. 2009). *K* values were varied from 2 to 8. For each *K* value, 10 replicate ADMIXTURE runs were performed using seeds from 1 to 10. Post-processing and visualization of ADMIXTURE results were carried out using the CLUMPAK web server (Kopelman et al. 2015). The major modes identified by CLUMPAK are presented. Results from *K* > 3 appear no longer informative and are not shown. SNVs and MNVs in non-intronic sequences were visualized in a heatmap by employing the R package ComplexHeatmap. For the 29 types of *K*-region sequences, bi-allelic SNVs and MNVs were identified in the same way, and were used for heatmap analysis.

### Recombination analysis

A gap-stripped and intron-free alignment of 69 MT types was used as input for the RDP4 program (Martin et al. 2015). Seven statistical methods implemented in RDP4, including RDP, GENECONV, BootScan, Maxchi, Chimaera, SiScan, and 3Seq, were used for the detection of recombination events.

### Divergence time estimation

The *S. pombe* nuclear genome is mostly euchromatic, with heterochromatin only existing in centromeres, telomeres, mating-type region, and rDNA. Heterochromatic regions tend to have higher mutation rates (Polak et al. 2015; Sun et al. 2016). Because the *K*-region, being part of the mating-type region, is heterochromatic (Grewal & Klar 1997), it may have a mutation rate higher than the mutation rate of the nuclear genome as a whole. To obtain a calibrated mutation rate for the *K*-region, we chose two strains reflecting ancestral genetic divergence (Tusso et al. 2019): the pure *Sp* lineage strain JB869 (*Sp* ancestry proportion 0.98) harbouring the K1 type *K*-region sequence and the pure *Sk* lineage strain JB758 (*Sk* ancestry proportion 0.98) harbouring the K19 type *K*-region sequence. We performed the calibration by comparing the SNV differences between JB869 and JB758 in the *K*-region versus the SNV differences between these two strains in the feature-free regions of the nuclear genome, which lack any PomBase-annotated features including genes and other genomic features such as repeat regions and low-complexity regions (Lock et al. 2019). We chose the feature-free regions for comparison because the *K*-region lacks protein-coding genes and is probably under little selective constraint and because the feature-free regions are also subject to little purifying selection (Jeffares et al. 2015). The coordinates of the feature-free regions were extracted from the genome annotation files chromosome1.contig, chromosome2.contig, and chromosome3.contig downloaded from ftp://ftp.pombase.org/pombe/genome_sequence_and_features/artemis_files/ (last modification dates of these annotation files are all 2017/10/18). Using published variant data (Jeffares et al. 2015), we found that JB869 and JB758 differ by 70 SNVs in the 6.53-kb *K*-region and 12,321 SNVs in the 1.465-Mb feature-free regions. Thus, the SNV density of the *K*-region was 27% higher than that of the feature-free regions of the nuclear genome. Essentially the same difference in SNV density was obtained when making comparisons using other pure-lineage *Sp* and *Sk* strains (Tusso et al. 2019). We interpret this difference in SNV density as a 27% increase in the mutation rate of the *K*-region relative to the mutation rate of the remaining nuclear genome. Using 1.27 as the calibration parameter and 2.00 × 10^−10^ mutations per site per generation as the genome-wide mutation rate (Behringer & Hall 2015; Farlow et al. 2015), we calculated the mutation rate of the *K*-region to be 2.54 × 10^−10^ mutations per site per generation. BEAST version 2.4.7 and its associated programs were used for divergence time estimation (Bouckaert et al. 2014). All positions in the *K*-region were used. The site model was selected using bModelTest version 1.0.4 and assuming a strict molecular clock. We compared two tree priors, the Yule model and the birth-death model, and chose the latter according to evaluations performed using Tracer version 1.6. Five independent runs were performed for 10 million generations each. We initiated runs on random starting trees, and sampled the trees every 10,000th generation. Effective sampling sizes were above 200 for all parameters. Results of the five runs were combined, with 10% removed as burn-in, using LogCombiner. Maximum clade credibility trees were summarized using TreeAnnotator, with posterior probability limit set to 0.5. Trees were visualized using FigTree.

## RESULTS

### *De novo* assembly of the complete mitogenomes of 192 *S. pombe* strains

A previous study generated Illumina-based genome sequencing data of 161 *S. pombe* isolates (JB strains) (Jeffares et al. 2015). Based on SNVs in the nuclear genome, these JB strains were found to have 57 types of nuclear genomes, with 129 strains falling into 25 “clonal clusters” each composed of multiple strains with near-identical nuclear genomes and 32 other strains each possessing a uniquely distinct nuclear genome. The authors of that study chose a set of 57 isolates, called “non-clonal strains”, to represent the 57 types of distinct nuclear genomes that differ from each other by no less than 1,900 SNVs (Jeffares et al. 2015). We used the publicly available Illumina sequencing data of these 161 JB strains (Supplementary Tables S1 and S2) to perform *de novo* assembly of their mitogenomes. We were able to assemble the complete circular-mapping mitogenomes of 154 (96%) JB strains (Supplementary Table S1). Validation using PacBio- and MinION-generated long read data of 15 diverse JB strains (Tusso et al. 2019) shows no difference to the Illumina-based mitogenome assemblies, corroborating their accuracy. The 154 assembled mitogenomes encompass 59 non-identical sequence types, which we term MT types (Supplementary Table S1). MT types may differ by as little as one nucleotide.

The 59 MT types present among the JB strains by and large correlated with the 57 previously defined nuclear genome types present among these strains (Supplementary Table S1). 55 of the 57 non-clonal strains have fully assembled mitogenomes. These 55 mitogenomes fall into 53 MT types, with three non-clonal strains, JB1205, JB1206, and JB1207, sharing the same MT type. The two non-clonal strains without fully assembled mitogenomes, JB842 and JB874, belong to clonal clusters 18 and 24, respectively. Other strains belonging to these two clusters do have fully assembled mitogenomes, which indicate that these two clusters correspond to two additional MT types (MT47 and MT21). The remaining 4 MT types (MT25, MT39, MT52, and MT65) are each highly similar to the MT type of a non-clonal strain and thus represent intra-cluster variations, with differences being a single SNV (MT25 vs. MT26 for cluster 10 and MT52 vs. MT53 for cluster 2), a single one-nucleotide indel (MT65 vs. MT64 for cluster 23), or the presence-absence polymorphisms of mitochondrial introns (MT39 vs. MT40 for cluster 15). The fact that we have obtained full-length mitogenomes from JB strains representing all 57 nuclear genome types indicates that the mitogenome diversity among the JB strains has been comprehensively captured.

To explore intraspecific diversity beyond that of the JB strains, we obtained 38 additional *S. pombe* isolates (DY strains) from Chinese and US culture collections, performed genome sequencing on them, and assembled the full-length mitochondrial genomes for all of them (Supplementary Table S1). The resulting 38 mitogenomes fall into 19 non-identical types, including 9 MT types present among the JB strains, and 10 MT types not present among the JB strains. Thus, overall, we identified 69 MT types from the fully assembled mitogenomes of 192 *S. pombe* isolates. We performed gene annotation on these MT types. The annotated sequences of the 69 MT types have been deposited at GenBank (accession numbers MK618072–MK618140).

### Identification of 20 errors in the reference *S. pombe* mitogenome

The reference *S. pombe* mitogenome (accession numbers NC_001326.1 and X54421.1) was from a Leupold-background strain with the genotype *h^−^ ade7-50* (Lang 1984; Zimmer et al. 1984). The reference *S. pombe* nuclear genome was derived from Leupold’s 972 *h^−S^* strain (Wood et al. 2002), called JB22 in the JB strain set. JB22 is the non-clonal strain representing clonal cluster 1 of the JB strains (Jeffares et al. 2015, 2017). Our *de novo* assembly of the mitogenomes showed that the mitogenomes of all clonal cluster 1 JB strains are identical. We designate this type of mitogenome MT1 (accession number MK618072).

Despite both being from the Leupold strain background, MT1 differs from the reference *S. pombe* mitogenome in 20 positions, including 13 single-nucleotide substitutions, 1 double-nucleotide substitution, 5 single-nucleotide indels, and 1 three-nucleotide insertion (Table 1). 13 of these 20 differences are located in protein-coding genes, and 11 of them alter amino acid sequences (Table 1). A previous study has uncovered 16 of these 20 differences by mapping Illumina sequencing reads to the reference mitogenome, but did not ascertain whether the differences were due to reference errors or polymorphisms (Iben et al. 2011). Our mitogenome assemblies show for all of these 20 positions that the sequence of MT1 is identical to those of the other 68 MT types. Thus, these 20 differences are caused by reference errors, not by naturally existing polymorphisms.

### Discovery of two new mitochondrial introns

Using restriction fragment analysis, a previous study of 26 *S. pombe* isolates estimated that the mitogenomes vary in length from 17.6 to 24.6 kb (Zimmer et al. 1987). We found here that among the 69 MT types, the mitogenome length varied between 17618 bp and 26910 bp. Length variation is almost entirely due to intron presence-absence polymorphisms (Figure 1A and Supplementary Table S3; intron presence-absence polymorphisms are described in more detail in a later section).

**Figure 1.**
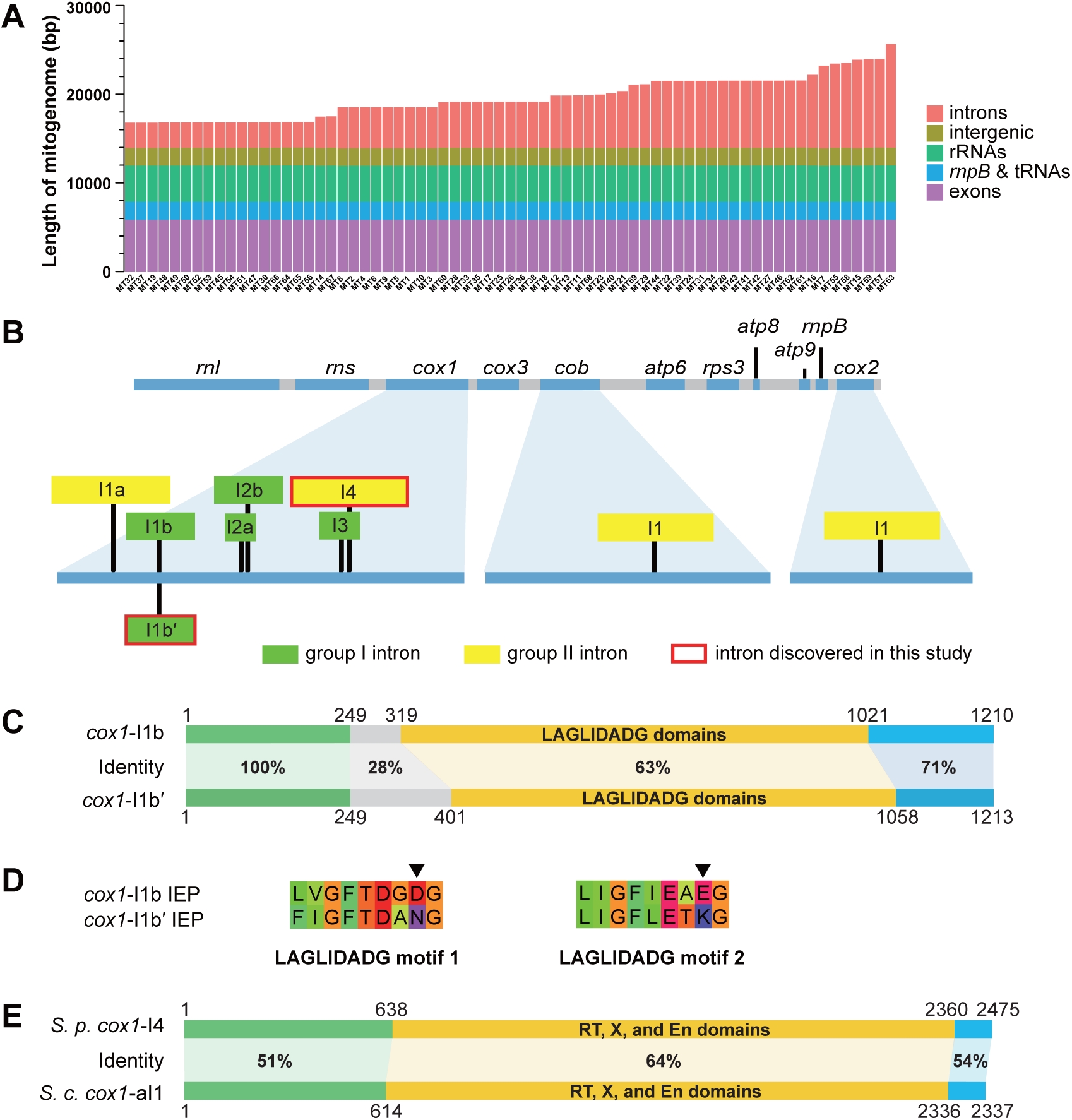
Length polymorphism among 69 types of *S. pombe* mitogenomes (MT types) and two newly discovered mitochondrial introns. (A) Length variation among 69 MT types ordered from the shortest to the longest. The sequence of each MT type is divided into 5 feature categories. The total length of each category is shown here in colour and also listed in Supplementary Table S3. (B) Diagram depicting the locations of the 9 distinct *S. pombe* mitochondrial introns. At the top, an intron-less mitogenome is depicted, with the lengths of genes and intergenic sequences drawn to scale. At the bottom, *cox1*, *cob*, and *cox2*, the three genes harbouring introns, are enlarged, and intron insertion positions are denoted by black vertical lines. Introns are shown as rectangles. The lengths of introns are not drawn to scale. (C) Comparison between the *cox1*-I1b intron in MT1 and the *cox1*-I1b′ intron in MT53. The DNA sequences of these two introns are divided into four segments: a segment coding for the LAGLIDADG domains, two segments upstream of the LAGLIDADG segment (one with 100% identity between *cox1*-I1b and *cox1*-I1b′ and the other with much lower identity), and a segment downstream of the LAGLIDADG segment. For each segment, percentage sequence identity (based on the length of sequence alignment) is shown. (D) The amino acid sequences of the two LAGLIDADG motifs in the intron-encoded proteins (IEPs) of *cox1*-I1b and *cox1*-I1b′. In each motif, the 8th residue critical for endonuclease activity is denoted by an arrowhead. (E) Comparison between the *cox1*-I4 intron in MT11 and the *cox1*-aI1 intron in the reference *Saccharomyces cerevisiae* mitogenome (NC_001224.1). The DNA sequences of these two introns are divided into three segments: a segment coding for the RT domain (reverse transcriptase domain), X domain (maturase domain), and En domain (endonuclease domain, formerly known as Zn domain), a segment upstream of the RT-X-En segment, and a segment downstream of the RT-X-En segment.

In *S. pombe,* there are seven previously known mitochondrial introns, which are called *cox1*-I1a, *cox1*-I1b, *cox1*-I2a, *cox1*-I2b, *cox1*-I3, *cob*-I1, and *cox2*-I1 (Schäfer 2003) (Figure 1B). Three of them, *cox1*-I1b (Schäfer et al. 1991), *cox1*-I2b (Lang 1984), and *cob*-I1 (Lang et al. 1985), are present in the Leupold strain. The other four introns, *cox1*-I1a (Schäfer & Wolf 1999), *cox1*-I2a (Trinkl & Wolf 1986), *cox1*-I3 (Trinkl & Wolf 1986), and *cox2*-I1 (Schäfer et al. 1998), are absent in the Leupold strain. In our *de novo* assembled mitogenomes, we identified two new introns, which we named *cox1*-I1b′ and *cox1*-I4, respectively (Figure 1B). *cox1*-I1b′ is located at the exact same position as *cox1*-I1b, and its presence is mutually exclusive with the presence of *cox1*-I1b. Both *cox1*-I1b′ and *cox1*-I1b are group I introns, and both encode proteins containing two LAGLIDADG endonuclease domains. The first 249 nucleotides of these two introns are identical, but the remaining portions are rather divergent, with the LAGLIDADG-domain-coding sequences exhibiting only 63% identity (Figure 1C). Despite this divergence, among the proteins in the NCBI non-redundant (nr) database, the closest homolog of the intron-encoded protein (IEP) of *cox1*-I1b′ is the IEP of *cox1*-I1b, suggesting recently shared ancestry of these two proteins (Supplementary Figure S1).

The IEP of *cox1*-I1b has been shown to possess both homing endonuclease and intron maturase activities (Pellenz et al. 2002; Schäfer 2003; Schäfer et al. 1994). For LAGLIDADG proteins, the nuclease activity requires that the 8th residue in the namesake LAGLIDADG motif must be an acidic residue to allow coordination with metal ions essential for catalysis (Chevalier et al. 2004). The 8th residues of the two LAGLIDADG motifs in the IEP of *cox1*-I1b are acidic residues (Figure 1D). In contrast, the 8th residues of the two LAGLIDADG motifs in the IEP of *cox1*-I1b′ are non-acidic residues (Figure 1D), suggesting that this protein probably has lost the homing endonuclease activity and acts solely as a maturase. This kind of degeneration of endonuclease function is remarkably common among the *Schizosaccharomyces* group I intron IEPs, as half of them (7/14) have non-acidic residues at the 8th position of at least one LAGLIDADG motif (Supplementary Figure S2).

In *S. pombe*, all previously analysed mitochondrial intron IEPs are thought to be translated as fusions with upstream exons, as the coding sequences of IEPs are always in-frame with 5′ exons (Schäfer 2003). We observed an exception to this rule in several *cox1*-I1b′ sequences. In three MT types (MT52, MT53, and MT66), the LAGLIDADG-domain-coding sequences in *cox1*-I1b′ are out-of-frame with 5′ exons due to a one-nucleotide insertion about 70 bp upstream of the LAGLIDADG domains (Supplementary Figure S3). This observation raises the possibility that *S. pombe* mitochondrial intron IEPs may not always be translated as in-frame extensions of the preceding exons.

The other intron newly identified in this study, *cox1*-I4, is located at a position downstream of all previously known *cox1* introns in *S. pombe* (Figure 1B). It is a group II intron. Our phylogenetic analysis showed that the IEP of *cox1*-I4 does not share a close relationship with any of the other *Schizosaccharomyces* group II intron IEPs (Supplementary Figure S4). Instead, it is most closely related to the IEPs encoded by *cox1*-ai1 and *cox1*-ai2 introns in *Saccharomyces cerevisiae* and *cox1*-ai1 introns in other species of the family Saccharomycetaceae (Figure 1E and Supplementary Figure S4). Thus, *S. pombe cox1*-I4 may have arisen through horizontal transfer from a Saccharomycetaceae species.

### Phylogenetic relationship of 69 MT types based on non-intronic sequences

Using the non-intronic sequences of nine genes (*cox1*, *cox3*, *cob*, *atp6*, *atp8*, *atp9*, *cox2*, *rnl*, and *rns*), which are conserved among the four *Schizosaccharomyces* species, we constructed a neighbour-joining tree of the 69 MT types (Figure 2, left). In this tree, the 69 MT types mostly fall into two highly distinct clades, with the single exception being MT15, locating at an intermediate position between the two clades. The same tree topology was obtained when we constructed a maximum likelihood tree using the non-intronic sequences of the above nine genes plus *rps3* and *rnpB* (Supplementary Figure S5). The smaller of the two clades contains MT1, the mitogenome in the Leupold strain background, from which the *S. pombe* reference genome was derived. Thus, we term this clade containing 14 MT types the REF clade. Accordingly, we term the other clade, which contains 54 MT types, the NONREF clade.

**Figure 2.**
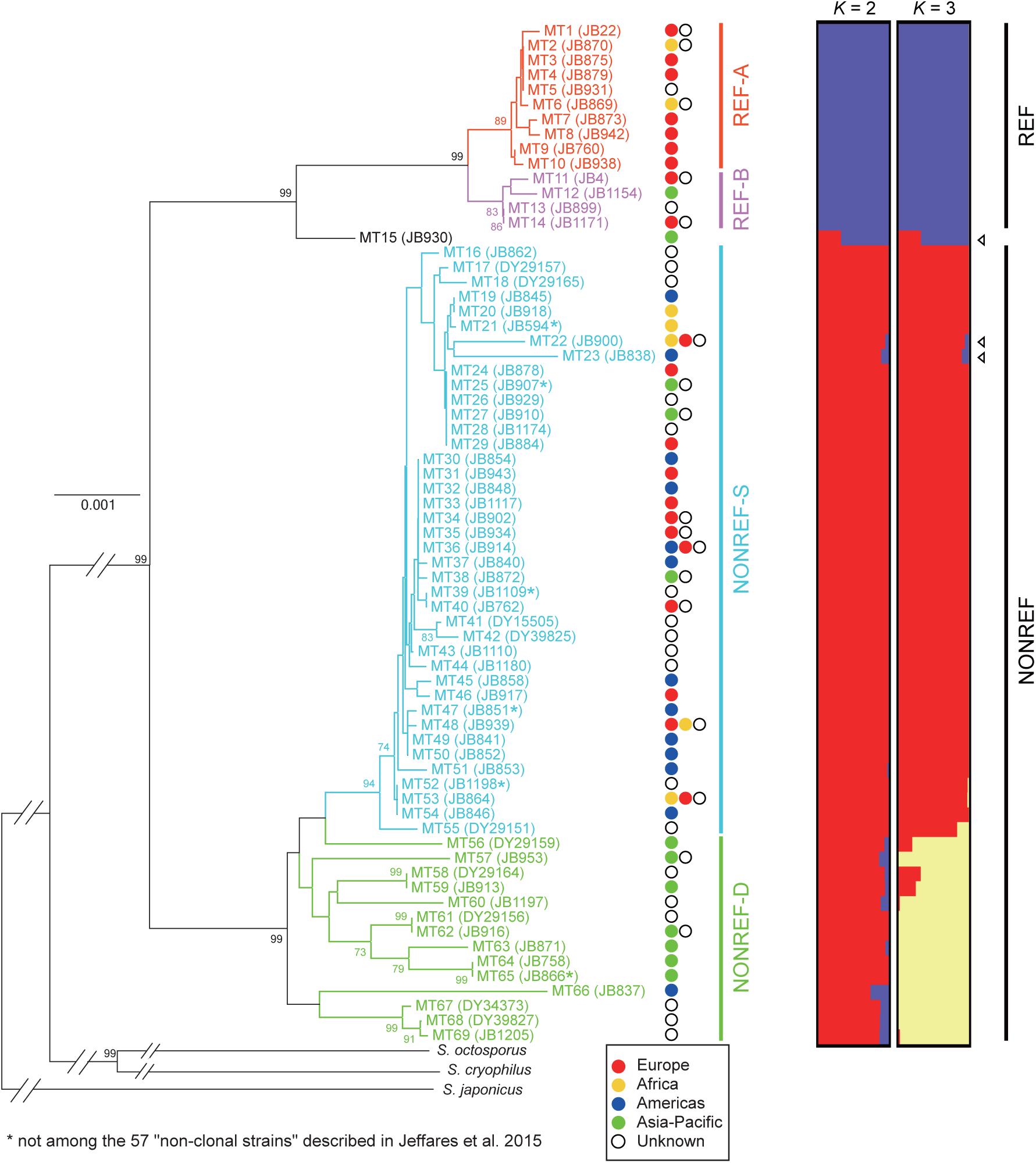
Phylogenetic analysis and maximum-likelihood clustering analysis of the non-intronic sequences of 69 MT types. For the phylogenetic analysis (left), a neighbour-joining tree was constructed using a concatenated alignment of the non-intronic sequences of 9 genes conserved across four *Schizosaccharomyces* species. The three non-*pombe Schizosaccharomyces* species (*S. octosporus*, *S. cryophilus*, and *S. japonicus*) were used as outgroup to root the tree. Outgroup branches are not drawn to scale. Bootstrap values higher than 70% are shown on the branches. Scale bar, 0.001 substitutions per site. For the maximum-likelihood clustering analysis using the ADMIXTURE program (right), the input was 222 segregating sites, which correspond to all bi-allelic SNV and MNV sites in non-intronic sequences (Supplementary Table S4). The 69 MT types, except for MT15, are classified into two clades (REF and NONREF clades), with each clade further divided into two subclades (A and B subclades for the REF clade, and S and D subclades for the NONREF clade). Arrowheads denote the three MT types that exhibit signs of inter-clade recombination. Tree branches and MT type names are coloured according to subclade affiliation. The name of a representative strain for each MT type is shown in parentheses. For the 59 MT types present among the JB strains, JB strains are chosen as representative strains, with preference given to the non-clonal strains. For each MT type, the continent(s) where associated strains have been isolated are indicated by coloured circles to the right of the tree, with the continent having more associated strains placed to the left (see Supplementary Table S1 for more detailed information on strains).

The REF clade has a substantially lower within-clade diversity than the NONREF clade. Nonetheless, MT types in the REF clade can be clearly divided into two subclades, which we term REF-A and REF-B. Within the NONREF clade, the relatedness among the MT types is highly uneven, with 40 MT types (MT16 to MT55) falling into a closely related monophyletic cluster, which we term NONREF-S subclade (S stands for similar). The large number of closely related MT types in the NONREF-S subclade may be partly due to non-random sampling of *S. pombe* isolates (see Discussion). The other 14 MT types (MT56 to MT69) in the NONREF clade are much more diverse, and we group them into a paraphyletic subclade, termed NONREF-D (D stands for diverse).

For the REF clade and the NONREF-D subclade, affiliated strains tend to share geographic origins (Figure 2, middle). MT types in the REF clade are mainly associated with strains collected from Europe, whereas MT types in the NONREF-D subclade are mainly associated with strains collected from Asia-Pacific. Among the 30 REF clade strains with known collection locations, 22 were collected from Mediterranean European countries including France, Spain, Italy, and Malta (Supplementary Table S1), suggesting a possible Southern European origin of this MT clade. Among the 8 NONREF-D strains with known collection locations, 6 were collected from Asian countries (Supplementary Table S1), suggesting that this subclade of high diversity is mainly distributed in Asia. In contrast, the NONREF-S subclade, despite its low internal diversity, has the broadest geographic distribution, with associated strains coming from all continents where *S. pombe* has been isolated. One possible explanation is that the NONREF-S strains have been distributed around the world by human migration (see Discussion).

To verify and complement the results obtained using phylogenetic tree construction, we identified from the alignment of non-intronic sequences a total of 222 bi-allelic SNVs and MNVs (Supplementary Table S4), and performed maximum-likelihood clustering analysis using the ADMIXTURE program (Figure 2, right) (Alexander et al. 2009). In addition, we directly visualized these bi-allelic SNVs and MNVs in a heatmap (Figure 3). These analyses lent support to the clade and subclade division. For the ADMIXTURE analysis, when *K* = 2, the REF clade and the NONREF clade are clearly distinguished; when *K* = 3, the NONREF clade is further separated into two clusters, corresponding to the NONREF-S subclade and the NONREF-D subclade. MT15, the MT type situated between the REF clade and the NONREF clade in the phylogenetic trees, exhibits an inter-clade mosaic pattern for both *K* values, suggesting that it may be a recombination product between REF and NONREF mitogenomes. Interestingly, two other MT types, MT22 and MT23, also consistently exhibit inter-clade mosaic patterns in the ADMIXTURE results, albeit to a lesser degree than MT15.

**Figure 3.**
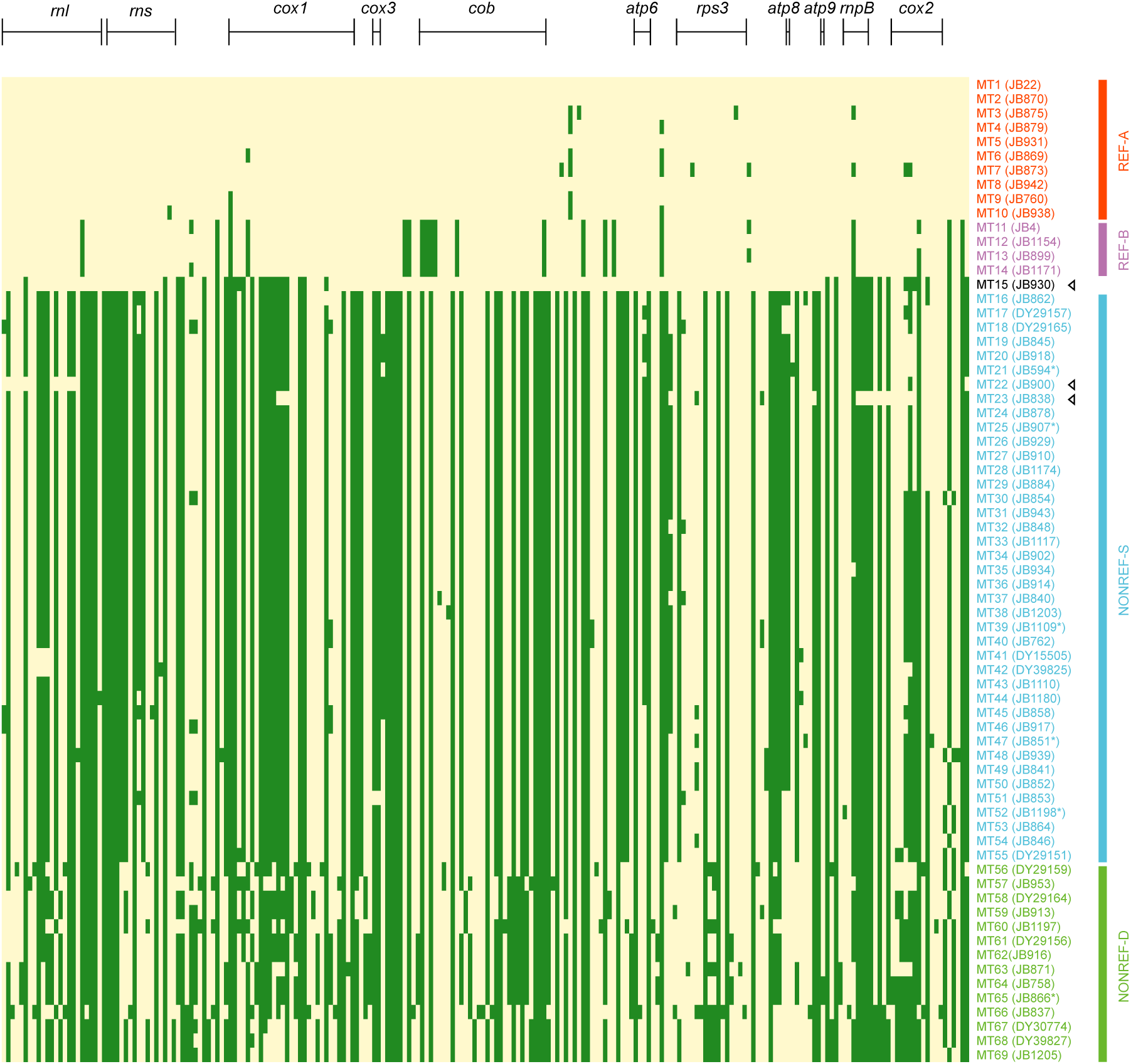
A two-colour heatmap of the 222 bi-allelic SNV and MNV sites in non-intronic sequences of 69 MT types. These are the same 222 sites used in the ADMIXTURE analysis shown in Figure 2. Each row in the heatmap represents an MT type (in the same order as in Figure 2), and each column represents a polymorphic site. MT1 alleles are coloured in yellow and non-MT1 alleles are coloured in green. Sites locating within the 8 protein-coding genes and the 3 large RNA genes are indicated by bracketed lines above the heatmap. MT type names and names of the representative strains are coloured according to subclade affiliation. Arrowheads denote the three MT types that exhibit signs of inter-clade recombination.

Inspecting the heatmap confirmed that MT15, MT22, and MT23 are the products of inter-clade recombination (Figure 3). MT15 appears to result from two inter-clade recombination events, with two stretches of its sequence resembling the REF-A subclade and the other two stretches resembling the NONREF-S subclade. The bulk of the sequences in MT22 and MT23 match those in other NONREF-S mitogenomes, with two small stretches of sequences in the *rnl* gene of MT22 and one small stretch of sequence spanning the *rnpB* gene in MT23 exhibiting REF clade patterns. We also performed statistical identification of recombination events using the program RDP4 (Martin et al. 2015) (Supplementary Table S5). Consistent with the results of ADMIXTURE analysis and visual inspection of the heatmap, the only recombination events supported by all seven recombination detection methods employed by RDP4 are those associated with MT15, MT22, and MT23. Recombination events not associated with these three MT types have much weaker support and appear to be mostly intra-clade recombination events occurring between NONREF mitogenomes.

Together, the above analyses of the non-intronic sequences demonstrate that present-day *S. pombe* mitogenomes descend from two well-separated ancient lineages, with only rare mitogenome recombination having occurred between lineages. This runs counter the the expectation from a previously published nuclear genome analysis concluding that *S. pombe* lacks strong population structure (Jeffares et al. 2015). The results are, however, consistent with an independent study published after the initial submission of this article revealing that *S. pombe* nuclear genomes also descend from two ancestral lineages (also see a later Results section) (Tusso et al. 2019).

### Presence-absence polymorphisms and phylogeny of mitochondrial introns

There are 18 types of intron presence-absence patterns in the 69 MT types (Figure 4A and Supplementary Table S3). For each of the 8 intron insertion sites, introns are only present in some but not all MT types, indicating that intron gain and/or loss have happened at all sites. The 4 group I intron sites are occupied in appreciably higher proportions (93%, 70%, 87%, and 67% for *cox1*-I1b/I1b′, *cox1*-I2a, *cox1*-I2b, and *cox1*-I3, respectively) than the 4 group II intron sites (41%, 14%, 28%, 54% for *cox1*-I1a, *cox1*-I4, *cob-*I1, and *cox2*-I1, respectively).

**Figure 4.**
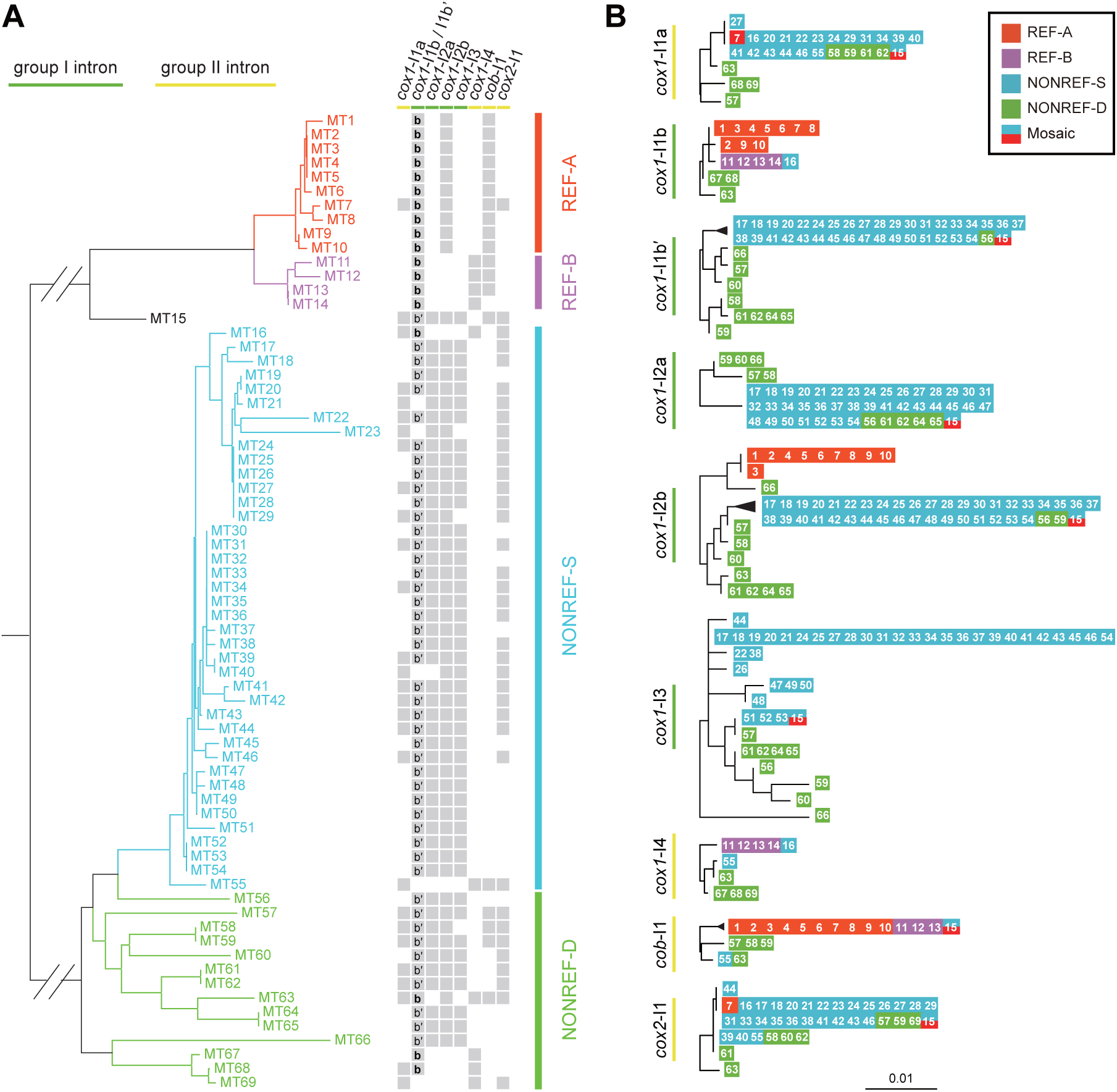
Intron presence-absence patterns and maximum likelihood trees of mitochondrial introns. (A) Intron presence-absence patterns at the 8 intron insertion sites in the 69 MT types. Grey squares indicate intron presence. The presence of *cox1*-I1b and *cox1*-I1b′ is respectively indicated by the bold letter b and the characters b′ inside a square. The phylogenetic tree on the left is the same as in Figure 2 without the outgroup branches. (B) Maximum likelihood trees of the 9 introns. The MT type from which the intron sequence originated is represented by the respective number in a coloured square. The colour of the square denotes the subclade affiliation of the MT type. The mosaic MT type, MT15, which partly resembles the REF-A subclade and partly resembles the NONREF-S subclade, is represented by a square coloured half-and-half with the colours of those two subclades. Trees were rooted by midpoint rooting. Scale bar, 0.01 substitutions per site.

The REF clade and the NONREF clade show distinct intron presence-absence patterns, with 6 of 8 intron sites exhibiting statistically significant differences between the clades (Supplementary Figure S6). *cox1*-I1a, *cox1*-I2a, *cox1*-I3, and *cox2*-I1 are completely or almost completely absent in the REF clade, but are common or ubiquitous in the NONREF clade. In contrast, *cob-*I1 is present in 93% of MT types in the REF clade but is present in only 9% of the MT types in the NONREF clade. For the *cox1*-I1b/I1b′ site, *cox1*-I1b is present in all MT types in the REF clade, whereas *cox1*-I1b′ is present in 83% of the MT types in the NONREF clade. These opposing patterns suggest that the two ancient *S. pombe* mitogenome lineages evolved different intron contents after their divergence.

Intron presence-absence patterns also exhibit a correlation with the subclade division within the REF clade. The REF-A subclade and the REF-B subclade are perfectly distinguished by the presence-absence patterns of *cox1*-I2b and *cox1*-I4, with the REF-A MT types all having *cox1*-I2b but not *cox1*-I4, and the REF-B MT types all having *cox1*-I4 but not *cox1*-I2b.

Within the low-nucleotide-diversity NONREF-S subclade, two group II introns, *cox1*-I1a and *cox2*-I1, are respectively present in 45% and 67.5% of the 40 MT types, and their presence-absence patterns do not obviously correlate with the nucleotide-based phylogeny, indicating that these two introns may have undergone extensive gain and/or loss events during the evolution of NONREF-S mitogenomes. Remarkably, all 18 NONREF-S MT types containing *cox1*-I1a also contain *cox2*-I1 (*P* = 0.00008, Fisher’s exact test), suggesting that, for reason(s) unclear to us, the presence of *cox1*-I1a in this subclade may be dependent on the presence of *cox2*-I1.

We constructed maximum likelihood trees for each of the 9 introns (Figure 4B). By and large, the phylogeny based on the sequences of a given intron mirrors the phylogeny of the MT types harbouring that intron, and shows a clear distinction between the REF clade and the NONREF clade for the four introns with appreciable presence in both clades (*cox1*-I1b, *cox1*-I2b, *cox1*-I4, and *cob-*I1). Thus, during *S. pombe* evolution, mitochondrial introns have rarely crossed the boundary between the two clades, consistent with the low extent of inter-clade recombination described earlier. There are a few notable exceptions. MT7 is the only REF clade MT type containing *cox1*-I1a and *cox2*-I1, and these two introns in MT7 are respectively identical to those in the majority of the NONREF-S MT types, suggesting that they may originate from cross-clade transfer. MT16 is one of few NONREF MT types harbouring *cox1*-I1b and *cox1*-I4, and these two introns in MT16 are respectively identical to those in the REF-B MT types but different from those in the other NONREF MT types, suggesting that they may also result from cross-clade transfer.

### *De novo* assembly and phylogenetic analysis of the *K*-region

As the above results indicate, contemporary *S. pombe* mitogenomes descend from two distinct ancient lineages. Next, we addressed the question whether *S. pombe* nuclear genomes also share a similar evolutionary history. Because a previously published nuclear genome analysis suggested that interbreeding between populations has occurred during the evolution of *S. pombe* (Jeffares et al. 2015), and such interbreeding (admixture) is expected to cause nuclear genome recombination that can interfere with phylogenetic inference (Posada & Crandall 2002), we reasoned that a nuclear genome region where recombination is repressed may be better suited for deducing the phylogenetic history of the *S. pombe* nuclear genome. Based on this rationale, we chose to analyse the *K*-region in the nuclear genome (Grewal & Klar 1997), which is situated between two donor mating-type loci *mat2* and *mat3*, and is a known “cold spot” for both meiotic recombination and mitotic recombination (Egel 1984; Thon & Klar 1993). Using the genome sequencing data of the 199 *S. pombe* strains described above, we performed read mapping analysis and found that among these strains, 12 lack the *K*-region (Supplementary Table S1). For 150 of the 187 *K*-region-containing strains, we obtained by *de novo* assembly the complete sequences of the two unique sections of the *K*-region, and found that these *K*-region sequences belong to 29 non-identical sequence types, which we term *K*-region types (Figure 5A and Supplementary Table S1). *K*-region types may differ by as little as one nucleotide.

**Figure 5.**
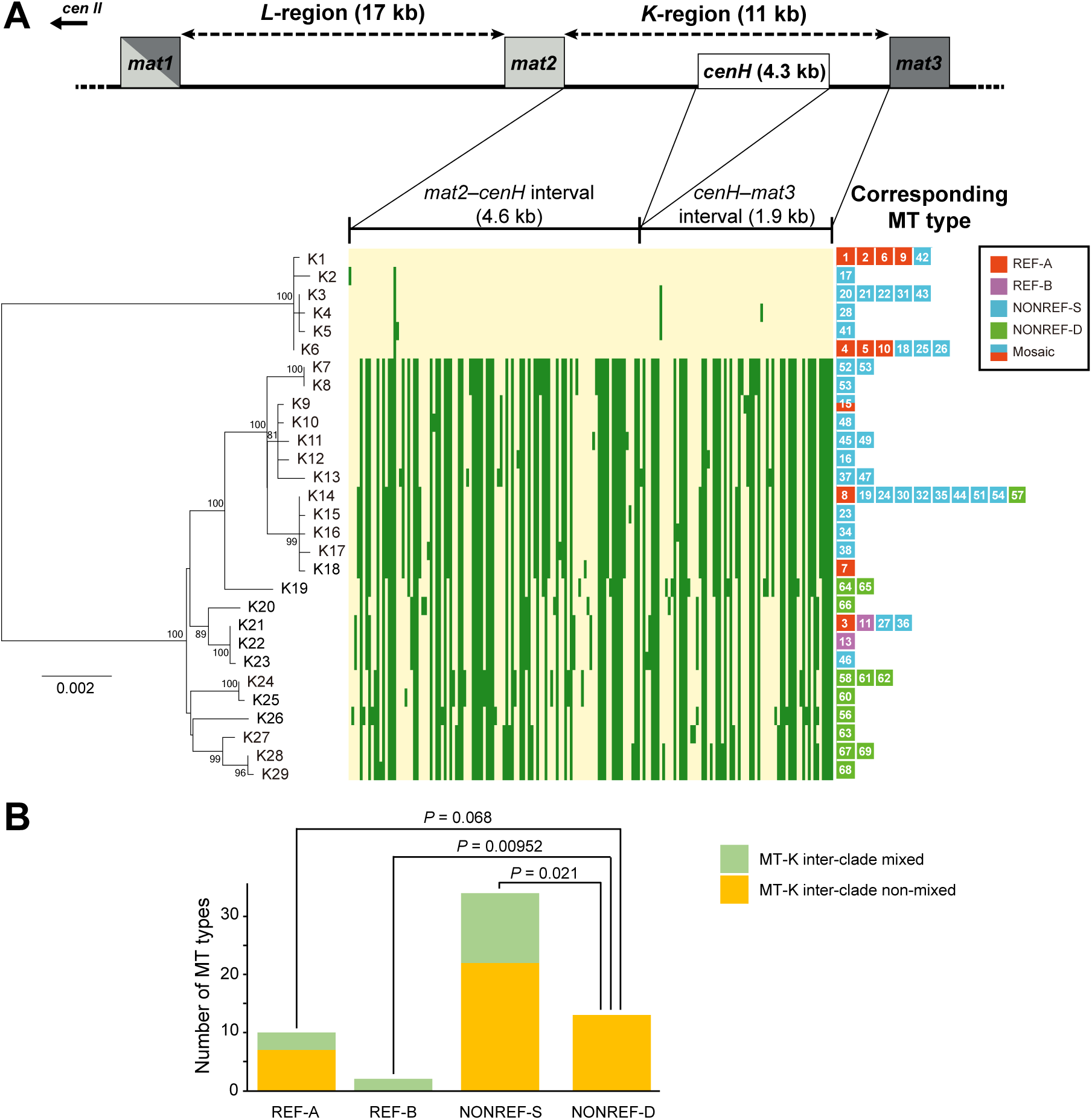
Phylogeny of the 29 *K*-region types and their relationship to MT types. (A) The part of chromosome II where the mating-type loci are located is depicted in a diagram at the top. The lengths of the *L*-region, the *K*-region, and the three *mat* genes are not drawn completely to scale. *cenH* is a centromere-repeat-like sequence that cannot be assembled from Illumina sequencing data. A neighbour-joining tree of the 29 *K*-region types (bottom left) was constructed. The tree was rooted by midpoint rooting. Bootstrap values higher than 70% are displayed on the tree. Scale bar, 0.002 substitutions per site. We also constructed a maximum likelihood tree which shows the same topology (Supplementary Figure S7). A heatmap (bottom middle) is used to visualize the 173 bi-allelic SNVs and MNVs in the *K*-region. K1 alleles are coloured in yellow and non-K1 alleles are coloured in green. MT types corresponding to each *K*-region type are displayed to the right of the heatmap. (B) If an MT type and its corresponding *K*-region type both belong to the low-diversity clade (group) or both belong to the high-diversity clade (group), the MT type is deemed “MT-K inter-clade non-mixed”. Otherwise, the MT type is deemed “MT-K inter-clade mixed”. For each MT type subclade, the numbers of MT types falling into these two categories are presented in a stacked bar chart. *P* values for between-subclade differences were calculated using Fisher’s exact test. Only *P* values < 0.1 are shown.

Phylogenetic analysis of the 29 *K*-region types shows that, like the mitogenomes, *K*-region sequences fall into two highly distinct clades (Figure 5A, bottom left, and Supplementary Figure S7). Moreover, similar to the situation in mitogenomes, the smaller clade, consisting of 6 *K*-region types (K1–K6), has a low internal diversity, whereas the larger clade, consisting of 23 *K*-region types (K7–K29), has a substantially higher internal diversity. For clarity, we refer to the two *K*-region clades as “groups”. The *K*-region type found in the reference nuclear genome, which we denote K1, belongs to the low diversity group. A heatmap analysis visualizing all bi-allelic SNVs and MNVs in the *K*-region sequences confirmed the deep divergence of the two groups and showed a lack of inter-group recombination (Figure 5A, bottom middle, and Supplementary Table S6). These results indicate that present-day *S. pombe* nuclear genomes also descend from two long-separated lineages, which probably correspond to the two ancient lineages of the mitogenomes. Based on this idea, the low-diversity *K*-region group should correspond to the REF clade of the MT types, and the high-diversity *K*-region group should correspond to the NONREF clade of the MT types.

The 150 strains with assembled *K*-region sequences are associated with 60 of the 69 MT types (Figure 5A, bottom right, and Supplementary Table S1). Any given MT type usually corresponds to only one of the 29 *K*-region type, except for MT53, whose associated strains have two types of *K*-regions (K7 and K8, differing by one single-nucleotide indel). The correlation between mitogenome clade affiliation and *K*-region group affiliation is only barely statistically significant (*P* = 0.042, Fisher’s exact test), with 58% (7/12) of REF clade MT types corresponding to *K*-region types in the low-diversity group, and 74% (35/47) of the NONREF clade MT types corresponding to *K*-region types in the high-diversity group. It is likely that interbreeding between populations has resulted in “MT-K inter-clade mixed” strains, in which the mitogenome and the *K*-region from different ancient lineages are brought together by hybridization.

We separately examined the extent of MT-K inter-clade mixing for each subclade of the MT types (Figure 5B). For the REF-A, REF-B, and NONREF-S subclades, 30% (3/10), 100% (2/2), and 35% (12/34) of the MT types are respectively MT-K inter-clade mixed. However, for the NONREF-D subclade, none of the 13 MT types are MT-K inter-clade mixed. Thus, strains harbouring the NONREF-D mitogenomes appear to have historically undergone less cross-lineage interbreeding.

Tusso et al. (2019) have independently identified two ancestral *S. pombe* lineages by performing in-depth analysis of the nuclear genomes of the 161 JB strains. They named the two lineages *Sp* and *Sk*, respectively (Tusso et al. 2019). Overall, the *Sp* lineage corresponds to the REF mitogenome clade and the low-diversity *K*-region group defined in this study: all pure *Sp* lineage JB strains (*Sp* ancestry proportion > 0.9, a criterion for pure-lineage strain used in Tusso et al. (2019)) fall into the REF mitogenome clade and the low-diversity *K*-region group (Supplementary Figures S8 and S9). Conversely, the *Sk* lineage corresponds to the NONREF mitogenome clade and the high-diversity *K*-region group. Consistent with the MT-K inter-clade mixing patterns described above, a large majority of JB strains with NONREF-D mitogenomes are pure *Sk* lineage strains (*Sk* ancestry proportion > 0.9), whereas most JB strains with NONREF-S mitogenomes have mosaic nuclear genomes (*Sp* ancestry proportions falling between 0.2 and 0.9). The only pure-lineage JB strains with NONREF-S mitogenomes are pure *Sk* lineage JB strains belonging to “clonal cluster 2” defined in Jeffares et al. 2015 (i.e. strains with MT52 and MT53 mitogenomes) (Supplementary Figure S8). This clonal cluster includes the “non-clonal strain” JB864 (Jeffares et al. 2015), which corresponds to NCYC132, the strain used in the 1950s and 1960s by Murdoch Mitchison in his cell cycle research (Mitchison 1970), and the type strain of *S. pombe*, CBS356, which was originally isolated from arak mash and was received by CBS in 1922 from the Král collection, the world’s first culture collection established in 1890 (Vaughan-Martini & Martini 2011). Given the high relatedness of NONREF-S mitogenomes, it is plausible that they may have all originated in the relatively recent past from one pure-lineage ancestral strain whose only unadmixed descendants among currently available *S. pombe* isolates are clonal cluster 2 strains.

### Estimation of the divergence time of the two ancient lineages of *S. pombe*

To obtain an estimate of the divergence time of the two ancestral lineages, we employed the Bayesian evolutionary analysis software BEAST to perform divergence dating on the 29 *K*-region types (Figure 6). Mutation accumulation studies have estimated a nuclear mutation rate of 2.00 × 10^−10^ substitutions per site per generation in *S. pombe* (Behringer & Hall 2015; Farlow et al. 2015). Applying a calibration (see Materials and Methods) we obtained a mutation rate estimation of 2.54 × 10^−10^ substitutions per site per generation for the *K*-region. Using this value as the mutation rate prior to perform BEAST analysis, we obtained a divergence time of 31.3 million generations for the two ancient lineages.

**Figure 6.**
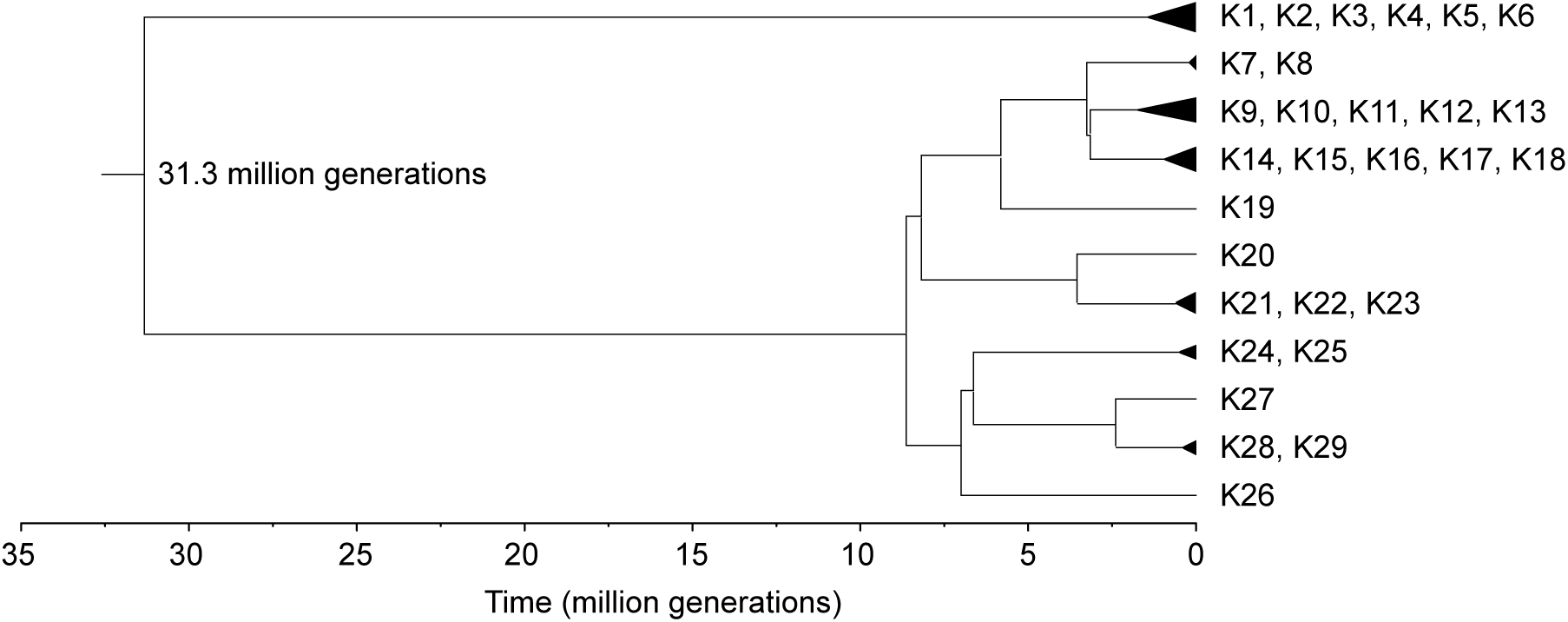
Divergence time of the two ancient lineages of *S. pombe* based on the *K*-region sequences was estimated using Bayesian evolutionary analysis implemented in BEAST 2.

## DISCUSSION

In this study, we *de novo* assembled and annotated full-length mitogenomes that encompass the mitogenome diversity of previously sequenced 161 JB strains (Jeffares et al. 2015) and 38 additional isolates (DY strains, this study). This comprehensive dataset allowed us to thoroughly examine the intraspecific mitogenome diversity existing among currently available *S. pombe* isolates and obtain new insights into the evolutionary history of this species.

Our analyses of the diversity patterns of mitogenome sequences and *K*-region sequences revealed that *S. pombe* isolates descend from two long-separated ancient lineages. The phenomenon of MT-K inter-clade mixing suggests that these two lineages have undergone admixture in recent historical time. The same conclusions have been reached in an independent study by Tusso et al. who analysed admixture proportions of the nuclear genome and the effect of admixture on phenotypic variation and reproductive isolation (Tusso et al. 2019). They revealed that a majority of the currently available *S. pombe* isolates have mosaic nuclear genomes that resulted from recent admixture between the two ancestral lineages.

Unlike in animals, where mitogenomes are usually inherited uniparentally, in fungi, biparental transmission of mitogenomes is common and thus allows the recombination between parental mitogenomes (Xu & Li 2015). For budding yeast species belonging to the family Saccharomycetaceae, naturally occurring mitogenome recombination appears to be common (Leducq et al. 2017; Peris et al. 2017; Wu & Hao 2014; Wu et al. 2015). In contrast, we show here that, despite a high level of inter-lineage admixture existing among the *S. pombe* isolates, inter-lineage recombination of *S. pombe* mitogenomes has rarely happened. A likely explanation is that, unlike the budding yeasts, *S. pombe* is a haplontic species, growing vegetatively as haploids, and only forming diploids transiently during sexual reproduction. The formation of a zygotic *S. pombe* diploid cell is immediately followed by meiosis and sporulation, and as a result, mitogenomes from two parental haploid cells may rarely have a chance to mix and recombine before being partitioned into four separate haploid progeny spores.

Even though *S. pombe* isolates have mostly been collected by chance rather than through dedicated search for this species, there have been a few cases of isolating multiple *S. pombe* strains from one relatively small geographic region. In particular, Carlos Augusto Rosa and his colleagues have isolated *S. pombe* from cachaça distilleries in the southeastern Brazilian state of Minas Gerais (Gomes et al. 2002; Pataro et al. 2000), and from the frozen fruit pulps acquired in markets in the eastern Brazilian state of Sergipe (Trindade et al. 2002). These Brazilian strains correspond to 10 MT types, with 6 MT types (MT45, MT47, MT49, MT50, MT51, and MT54) associated with the cachaça strains and 4 MT types (MT19, MT30, MT32, and MT37) associated with the fruit pulp strains. These 10 MT types all fall into the homogenous NONREF-S subclade and account for 25% of the MT types in this subclade, suggesting that non-random sampling partly contributes to the large size of this subclade.

In the 1960’s, Tommaso Castelli deposited into the DBVPG culture collection 13 *S. pombe* strains isolated from grape must and wine from the Mediterranean islands of Sicily and Malta (DBVPG online catalog, http://www.dbvpg.unipg.it/index.php/en/database). The six Sicily strains share the same MT type (MT9, subclade REF-A), whereas the seven Malta strains are associated with 3 MT types (MT4 in subclade REF-A, and MT24 and MT29 in subclade NONREF-S). The fact that MT types in both clades are found in strains isolated from wine-related substrates within a localized area (the size of Malta is only 316 km^2^) suggests ongoing opportunities for inter-clade exchange.

Given that the Rosa strains and the Castelli strains, the only notable *S. pombe* isolates with restricted geographic origins, together account for only 30% of the MT types in the NONREF-S subclade, the large size of this low-diversity subclade requires explanation(s) in addition to geographic sampling bias. Also in need of explanation are the extraordinarily wide distribution and the high extent of admixture of the strains associated with this subclade. We speculate that an ancestral pure-lineage *S. pombe* strain harbouring a NONREF-S mitogenome and a nuclear genome similar to that of JB864 may have by chance become associated with humans earlier than *S. pombe* strains harbouring other types of mitogenomes and, as a result, gained a world-wide distribution through co-migration with humans. In turn, the spreading of this strain may have also led to its encountering and hybridizing with REF clade strains. An alternative and non-exclusive explanation is that the NONREF-S mitogenomes may provide selective advantages in human-related substrates where *S. pombe* has most often been found, including cultivated fruits (raw and processed), cultivated sugar cane (raw and processed), and fermented beverages. We note that several JB strains with NONREF-D mitogenomes (JB913, JB1205, and JB1206) have mosaic nuclear genomes (Supplementary Figure S8), suggesting that ancestral pure-lineage strains harbouring NONREF-D mitogenomes have also contributed to inter-lineage admixture.

Based on our estimation, the two ancient lineages of *S. pombe* diverged about 31.3 million generations ago. To our knowledge, the shortest generation time (doubling time) reported for *S. pombe* under optimal laboratory growth conditions is approximately 2 hours (Johnson 1968). At such a growth rate, *S. pombe* can go through 12 generations per day, or 4,383 generations per year, and a divergence time of 31.3 million generations corresponds to 7,141 years. However, it is highly unlikely that *S. pombe* can proliferate continuously at this high rate in the wild. Taking inevitable encounters with unfavourable growth conditions into consideration, previous studies have estimated that the average generation time of *S. cerevisiae* in the wild can be more than 10 times longer than the shortest generation time observed in the laboratory (Fay & Benavides 2005; Ruderfer et al. 2006). Applying the same rationale, if we assume that *S. pombe* may go through as few as 400 generations per year in the wild, 31.3 million generations correspond to as many as 78,250 years. We emphasize that this is not a precise estimation of the divergence date because of the uncertainty on how to convert time from generations to years. Nevertheless, the divergence time of the two ancient lineages of *S. pombe* may fall within the most recent glacial period (“ice age”), which occurred from approximately 110,000 to 12,000 years ago (van Ommen 2015). The expansion of ice sheets and permafrost during a glacial period can lead to vicariance, the splitting of a population through the formation of geographic barriers (Hewitt 2000; Neiva et al. 2018). We speculate that glacial vicariance may have resulted in the allopatric separation of an ancestral population of *S. pombe* into isolated subpopulations. One of these subpopulations may have survived the glacial period in a glacial refugium in southern Europe, and become the low-diversity lineage with the REF clade mitogenomes.

It is of note that the mainly Asian distribution of the high-diversity pure-lineage NONREF strains (corresponding to the pure-lineage *Sk* strains described in Tusso et al. (2019)) is reminiscent of the situation of the other model yeast species *Saccharomyces cerevisiae*, whose highest intraspecific diversity exists in China (Wang et al. 2012). Based on this geographic pattern of diversity of *Saccharomyces cerevisiae* and other lines of evidence, the whole *Saccharomyces* species complex is now believed to have originated in Asia (Peter et al. 2018; Duan et al. 2018). It is possible that Asia is also a centre of origin of *S. pombe*.

The intraspecific *S. pombe* divergence patterns observed in this study are consistent with the following speculative evolutionary scenario: During the last glacial period, an ancient population of *S. pombe* was separated into refugia; one subpopulation suffered a bottleneck and became a low-diversity lineage mainly distributed in Southern Europe, whereas another subpopulation became a higher-diversity lineage mainly distributed in Asia; after the glacial period ended, perhaps aided by human migration, these two long-separated lineages came into secondary contact and began to hybridize; human migration has also shaped the worldwide distribution of *S. pombe*, and in particular, has spread strains with the NONREF-S mitogenomes to all over the world.

## AUTHOR CONTRIBUTIONS

Yu-Tian Tao performed the genome sequencing of the DY strains, analysed the assembled mitogenomes and *K*-region sequences, and prepared the manuscript; Fang Suo performed the *de novo* assembly of mitogenomes and *K*-region sequences; Yan-Kai Wang and Song Huang provided the Tn5 transposase; after the initial submission of this article, Sergio Tusso and Jochen Wolf performed mitogenome assembly validation using third-generation sequencing data, contributed the admixture proportion data, and edited the revised manuscript; Li-Lin Du devised and coordinated the project and together with Yu-Tian Tao wrote the manuscript.

## ACKNOWLEDGEMENTS

We thank Wen Hu for generating the Illumina sequencing library of DY15505. We thank Yang Liu and Wei Jiang for contributing to the mitogenome analysis at the early stage of this work. This work was supported by the Ministry of Science and Technology of China and by the Beijing Municipal Government.

