## Supplementary Figures for "Intraspecific diversity of fission yeast mitochondrial genomes"

Supplementary Tables S1–S6 are provided as a separate Excel file.

Supplementary Table S1. The 199 *S. pombe* isolates whose genome sequencing data are used in this study.

Supplementary Table S2. The 11 sequencing runs associated with wrong strain names.

Supplementary Table S3. The 69 MT types.

Supplementary Table S4. Bi-allelic SNVs and MNVs in the non-intronic sequences of the 69 MT types.

Supplementary Table S5. RDP4-detected recombination events among the mitogenomes.

Supplementary Table S6. Bi-allelic SNVs and MNVs in the K-region sequences of the 29 K-region types.

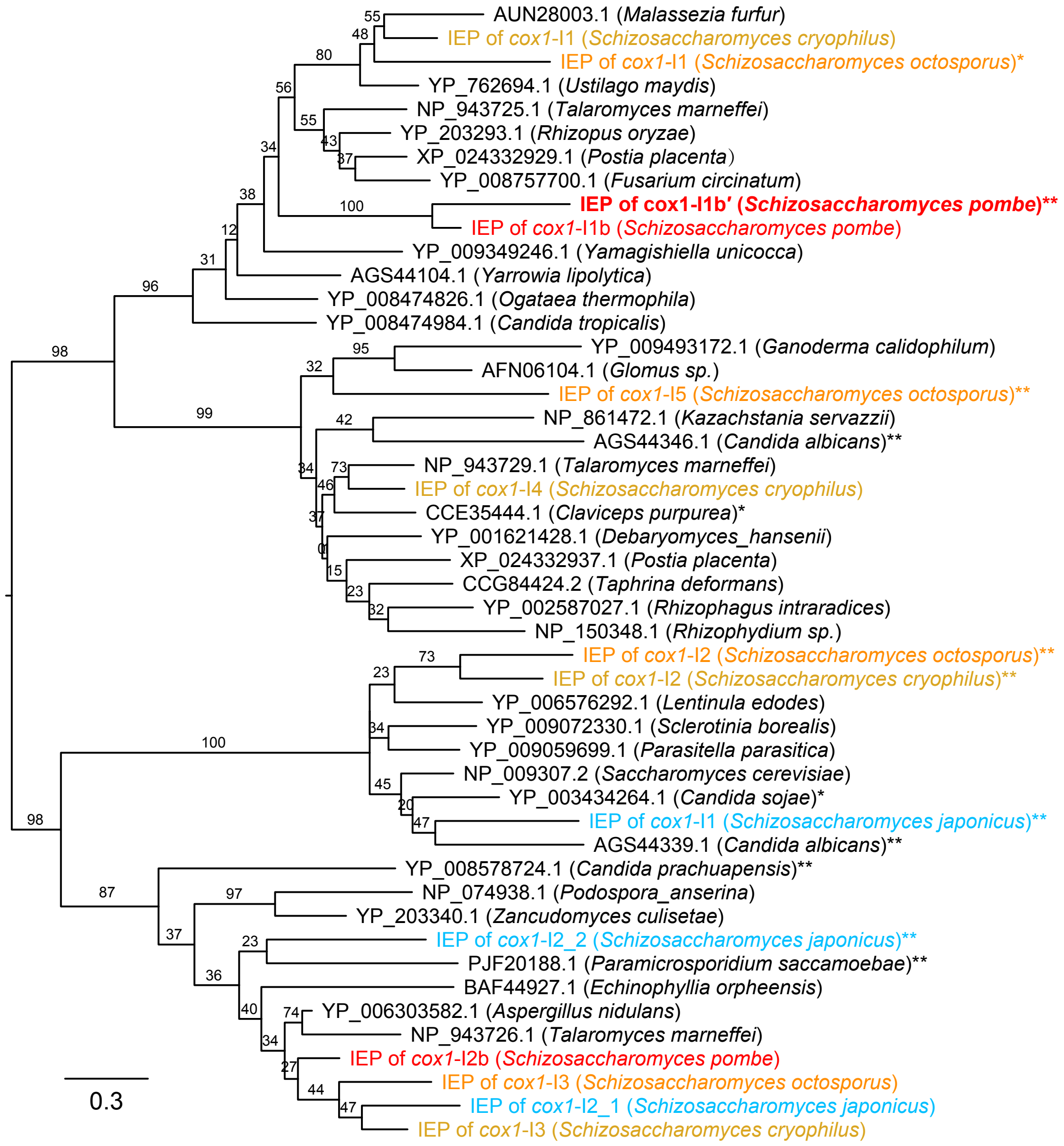

**Supplementary Figure S1**. Phylogenetic analysis of intron-encoded proteins (IEPs) of fission yeast group I introns and their close homologs. The IEPs encoded by group I introns in the mitogenomes of *S. pombe*, *S. octosporus*, *S. cryophilus*, and *S. japonicus* and their representative homologs retrieved by BLASTP search against the NCBI nr protein database are aligned using MAFFT. The sequence alignment of the conserved region, which corresponds to the two LAGLIDADG domains, is used for phylogenetic analysis. Maximum likelihood tree was constructed using MEGA. Tree was rooted by midpoint rooting. Bootstrap support values are shown on the tree branches. Scale bar, 0.3 amino acid substitutions per amino acid site. The names of

*Schizosaccharomyces* proteins are highlighted in colour (red for *S. pombe*, orange for *S. octosporus*, yellow for *S. cryophilus*, and blue for *S. japonicus*). The name of the IEP encoded by the newly discovered *S. pombe cox1*-I1b′ intron is highlighted in bold. Single asterisks indicate that the 8th residue in one of the two LAGLIDADG motifs is not an acidic residue. Double asterisks indicate that the 8th residue in neither LAGLIDADG motif is an acidic residue.

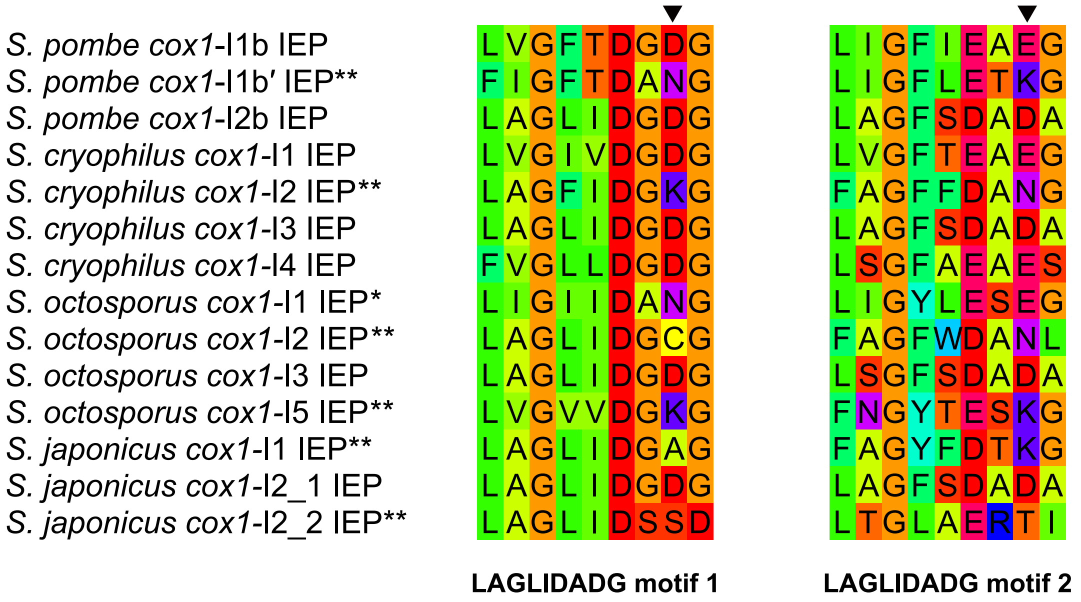

**Supplementary Figure S2**. The amino acid sequences of the two LAGLIDADG motifs in the IEPs of *Schizosaccharomyces* group I introns. In each motif, the 8th residue that needs to be an acidic residue to support the homing endonuclease activity is denoted by an arrowhead. Single asterisks indicate that the 8th residue in one of the two LAGLIDADG motifs is not an acidic residue. Double asterisks indicate that the 8th residue in neither LAGLIDADG motif is an acidic residue.

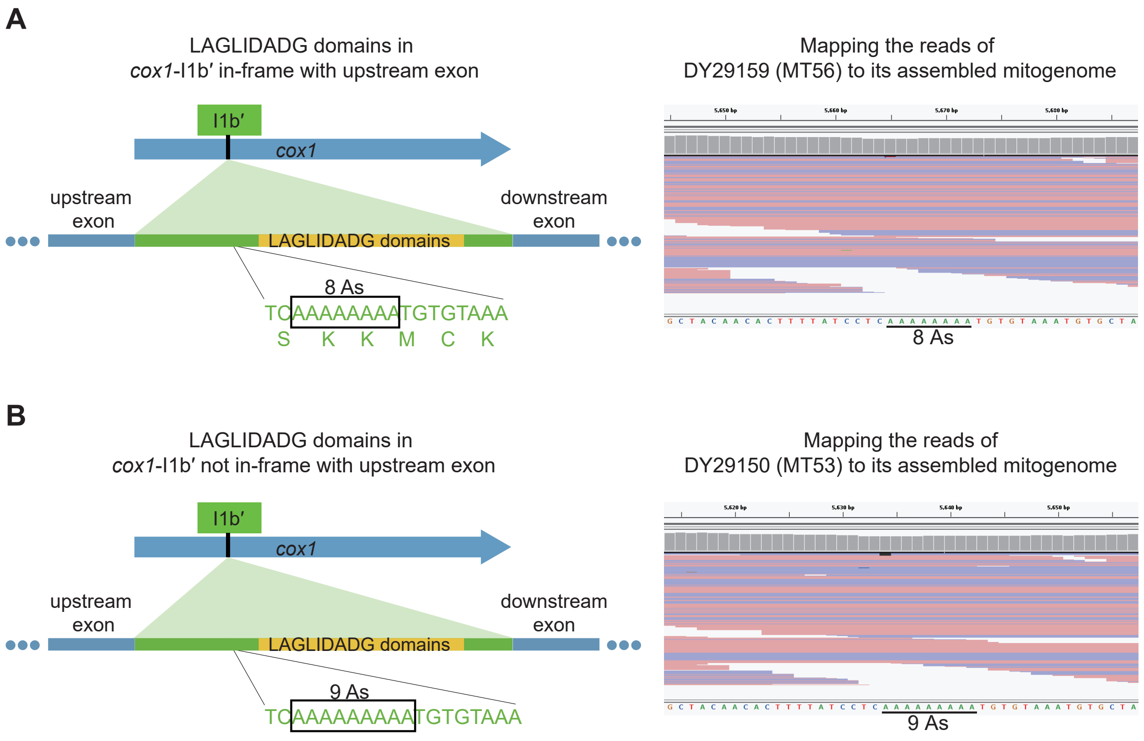

**Supplementary Figure S3**. The intron-encoded proteins (IEPs) in some *cox1*-I1b′ introns are not in frame with the upstream exons. A diagram depicting the location where a frameshift occurs in a polyA tract in some *cox1*-I1b′ introns (left) and an example of Illumina read mapping result (right) are shown for the in-frame situation (A) and the out-of-frame situation (B), respectively.

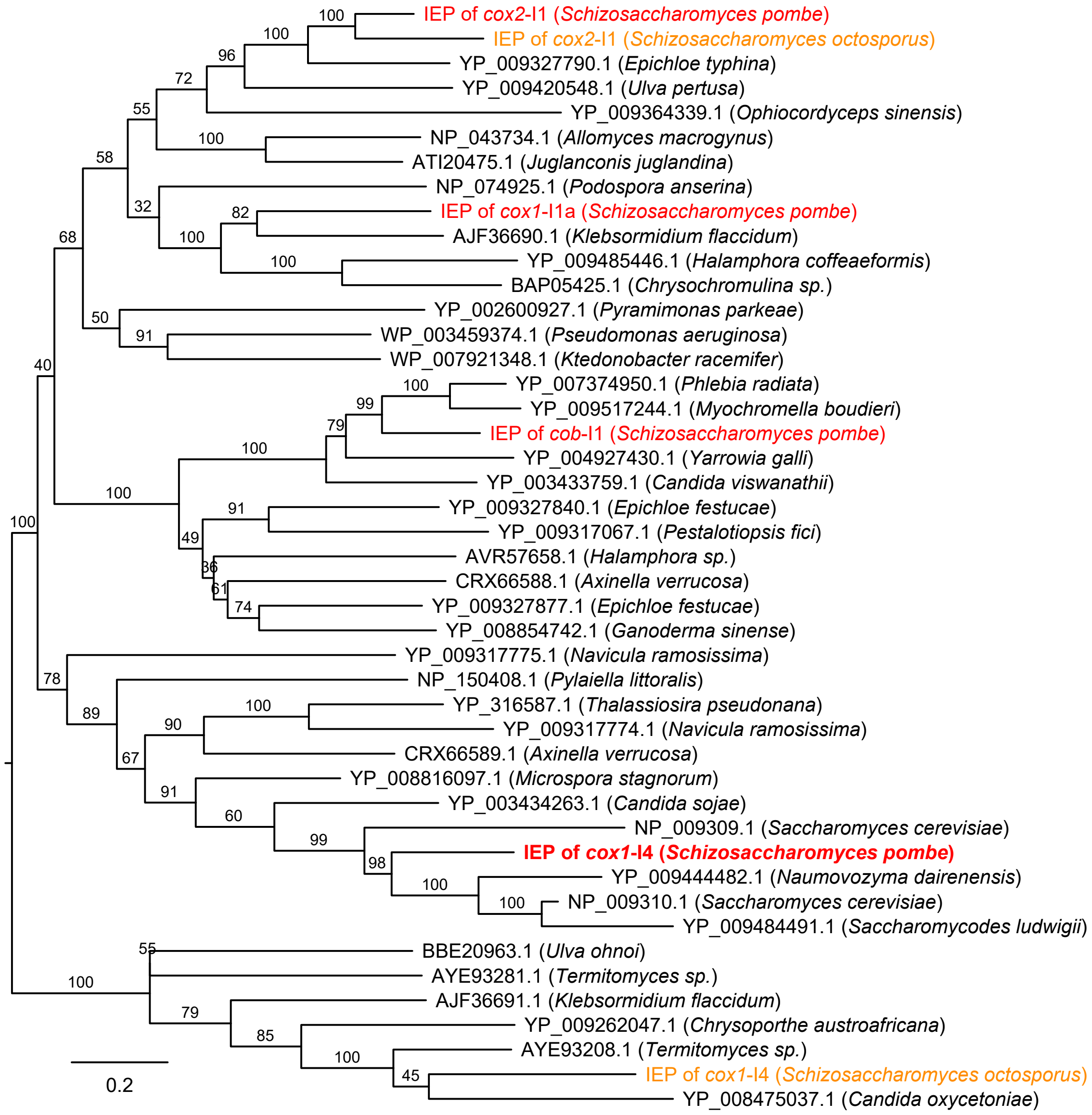

**Supplementary Figure S4**. Phylogenetic analysis of intron-encoded proteins (IEPs) of fission yeast group II introns and their close homologs. The IEPs encoded by group II introns in the mitogenomes of *S. pombe* and *S. octosporus* and their representative homologs retrieved by BLASTP search against the NCBI nr protein database are aligned using MAFFT (the mitogenomes of *S. cryophilus* and *S. japonicus* lack group II introns). The sequence alignment of the conserved region, which encompasses the RT domain, X domain, and En domain, is used for phylogenetic analysis. Maximum likelihood tree was constructed using MEGA. Tree was rooted by midpoint rooting. Bootstrap support values are shown on the tree branches.

Scale bar, 0.3 amino acid substitutions per amino acid site. The names of *Schizosaccharomyces* proteins are highlighted in colour (red for *S. pombe* and orange for *S. octosporus*). The name of the IEP encoded by the newly discovered *S. pombe cox1*-I4 intron is highlighted in bold.

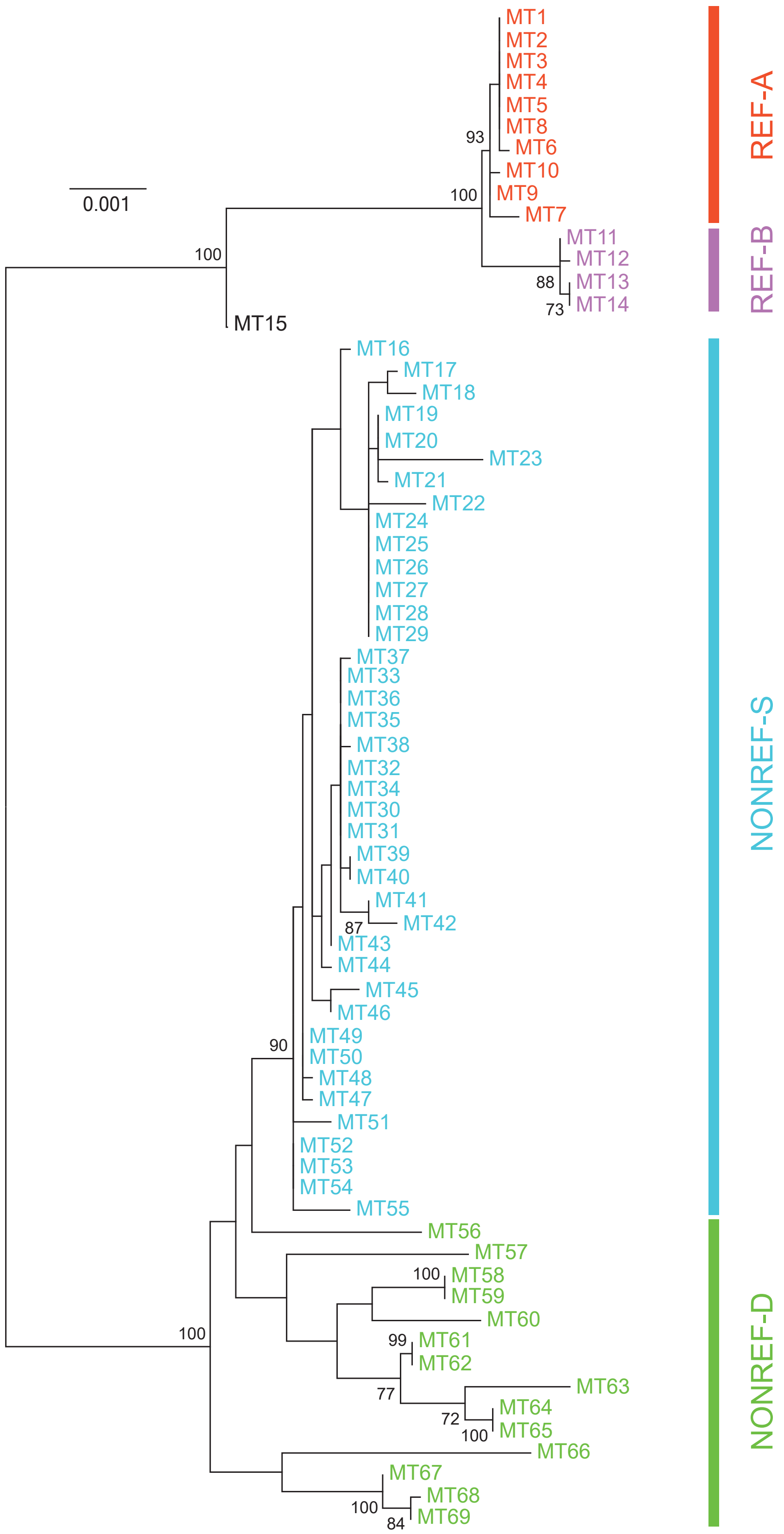

**Supplementary Figure S5**. Maximum likelihood tree constructed based on a concatenated alignment of the non-intronic sequences of 3 RNA genes (*rnl*, *rns*, and *rnpB*) and 8 protein-coding genes (*atp6*, *atp8*, *atp9*, *cob*, *cox1*, *cox2*, *cox3*, and *rps3*). Tree was rooted by midpoint rooting. Bootstrap values higher than 70% are displayed on the tree. Scale bar, 0.001 substitutions per site.

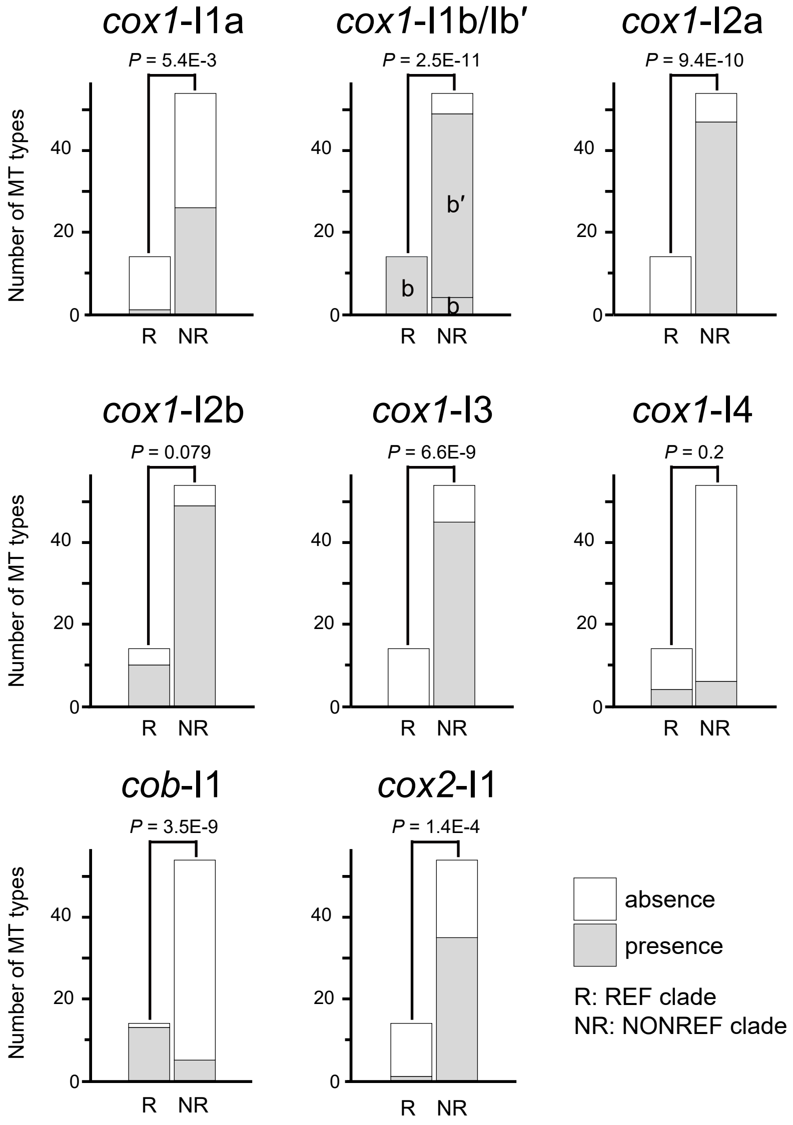

**Supplementary Figure S6**. Comparison between the REF clade and the NONREF clade for the intron presence-absence patterns at the 8 intron insertion sites. *P* values were calculated using Fisher's exact test.

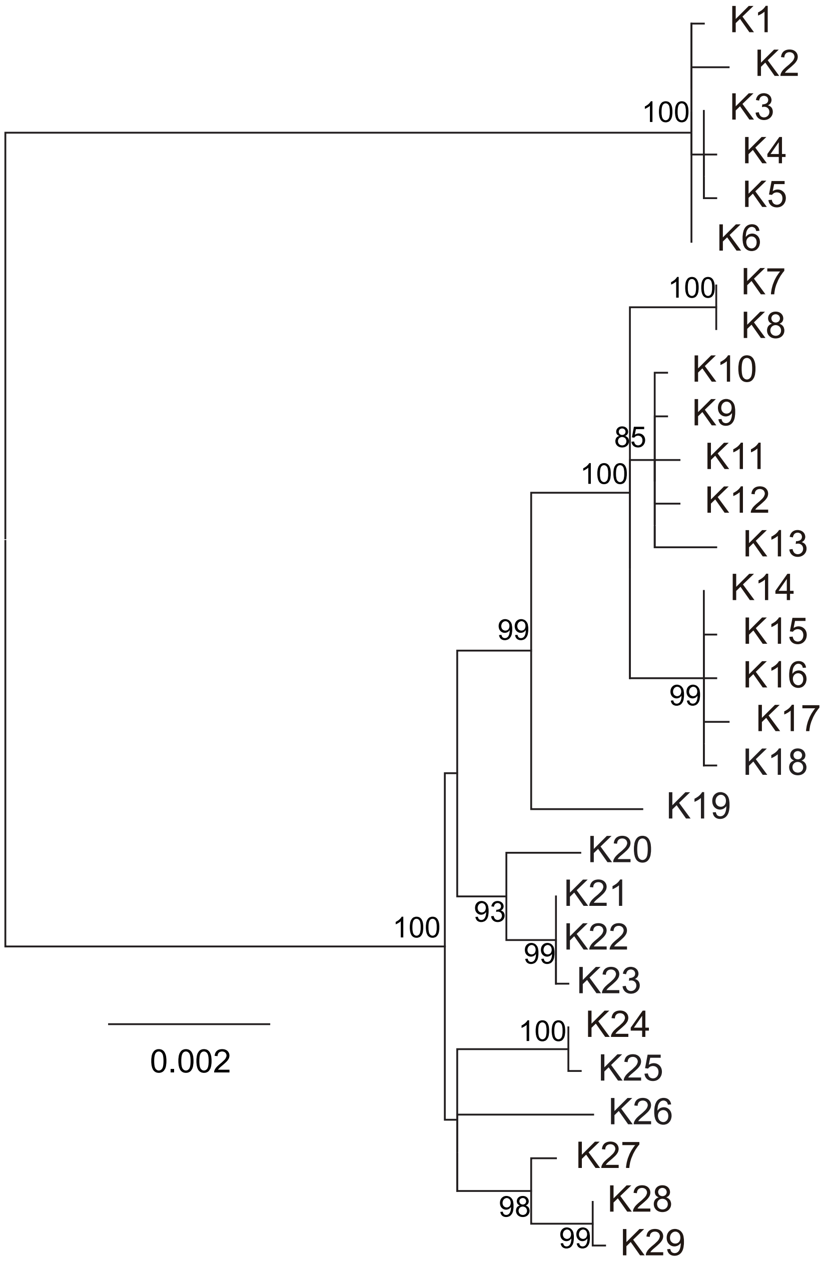

**Supplementary Figure S7**. Maximum likelihood tree constructed based on an alignment of *K*-region sequences. Tree was rooted by midpoint rooting. Bootstrap values higher than 70% are displayed on the tree. Scale bar, 0.002 substitutions per site.

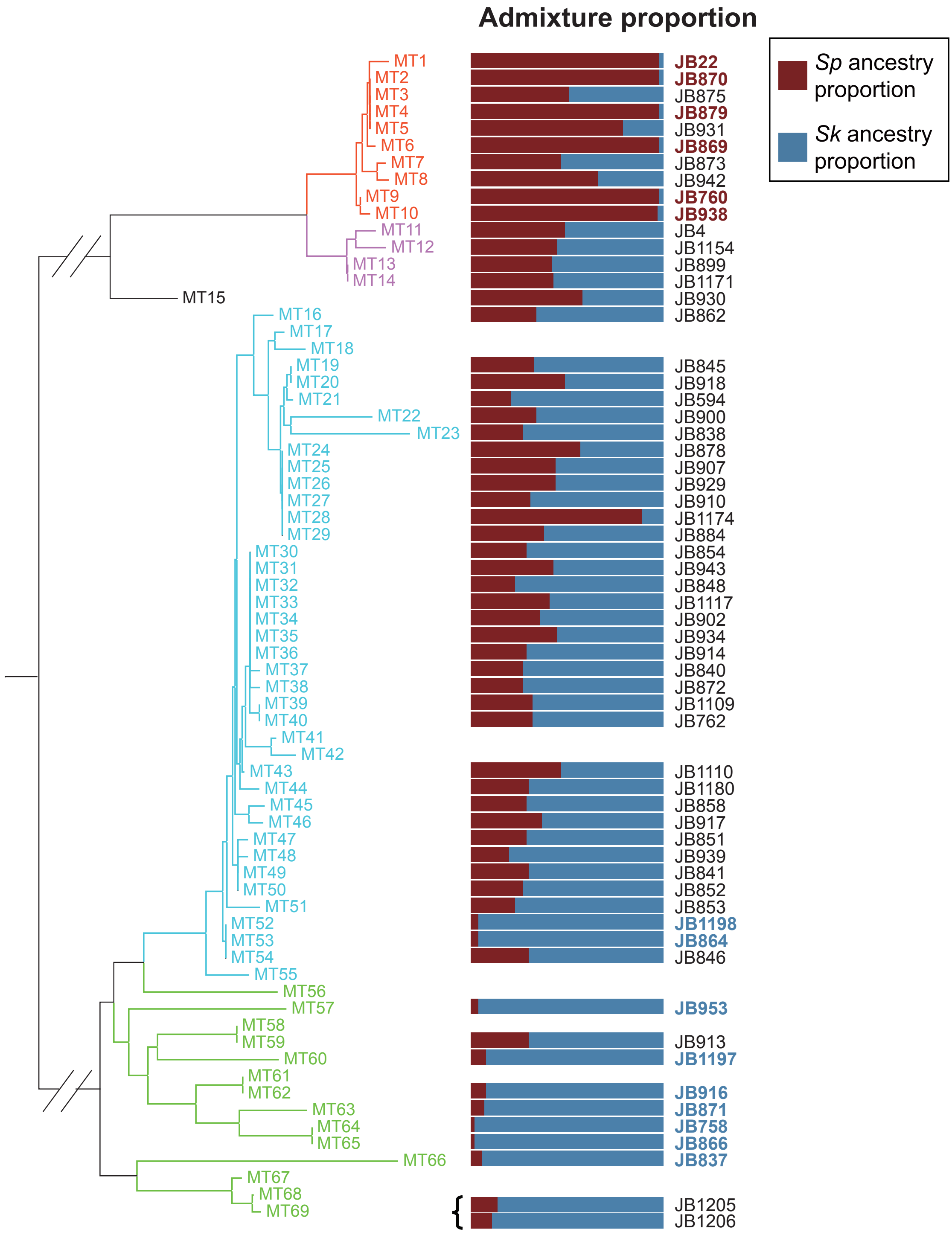

**Supplementary Figure S8**. The relationship between the MT types and the nuclear genome admixture proportions of the JB strains (proportions of the *Sp* lineage and the *Sk* lineage defined in Tusso et al. 2019). The phylogenetic tree on the left is the same one in Figure 2. Among the JB strains that have fully assembled mitogenomes, there are 59 MT types associated with 57 of the 57 nuclear genome types defined in Jeffares et al. 2015. For each of the 59 MT types, one representative JB strain for each associated nuclear genome type is shown, except for the nuclear genome type represented by JB1207, which is a diploid strain. When choosing the representative JB strain for a nuclear genome type, preference is given to the "non-clonal strains" defined in Jeffares et al. 2015. Pure-lineage strains (strains with *Sp* ancestry proportion or *Sk* ancestry proportion > 0.9) are highlighted by bold and colored font.

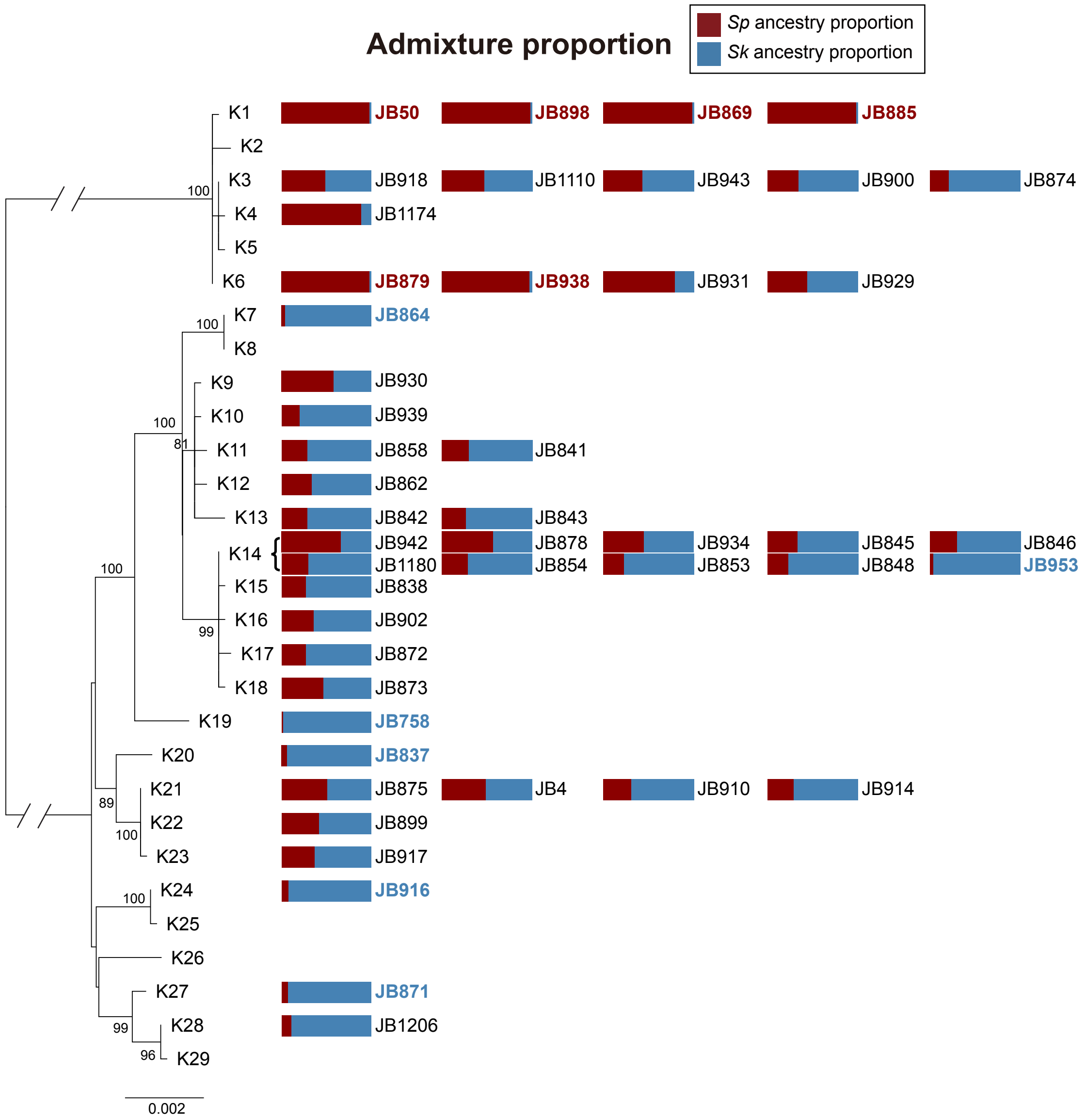

**Supplementary Figure S9**. The relationship between the *K*-region types and the nuclear genome admixture proportions of the JB strains (proportions of the *Sp* lineage and the *Sk* lineage defined in Tusso et al. 2019). The phylogenetic tree on the left is the same one in Figure 5A. Among the JB strains that have fully assembled *K*-region sequences, there are 23 *K*-region types associated with 47 of the 57 nuclear genome types defined in Jeffares et al. 2015. For each of the 23 *K*-region types, one representative JB strain for each associated nuclear genome type is shown. When choosing the representative JB strain for a nuclear genome type, preference is given to the "non-clonal strains" defined in Jeffares et al. 2015. Pure-lineage strains are highlighted by bold and colored font.
